# Regulators of male and female sexual development critical for transmission of a malaria parasite

**DOI:** 10.1101/2021.08.04.455056

**Authors:** Andrew J C Russell, Theo Sanderson, Ellen Bushell, Arthur M Talman, Burcu Anar, Gareth Girling, Mirjam Hunziker, Robyn S Kent, Tom Metcalf, Ruddy Montandon, Vikash Pandey, A Brett Roberts, Claire Sayers, Frank Schwach, Julian C Rayner, Thierry Voet, Katarzyna K Modrzynska, Andrew P. Waters, Mara K N Lawniczak, Oliver Billker

**Author notes:** These authors contributed equally to this work (Andrew J C Russell, Theo Sanderson, Ellen Bushell, Arthur M Talman). University of Vermont, Burlington VT 054051, USA. Wellcome Centre for Human Genetics, Roosevelt Dr, Headington, Oxford OX3 7BN, United Kingdom (Ruddy Montandon). Corresponding authors. For bulk transcriptomic time course and validation,; single cell analysis and validation,; functional screen and validation,.

## Abstract

The transmission of malaria parasites from vertebrate host to mosquito vector requires a developmental switch in asexually dividing blood-stage parasites to sexual reproduction. In *Plasmodium berghei* the transcription factor AP2-G is required and sufficient for this switch, but how a particular sex is determined in a haploid parasite remains unknown. Using a global screen of barcoded mutants, we here identify ten genes essential for the formation of either male or female sexual forms and validate their importance for transmission. High-resolution single-cell transcriptomics of wild-type and mutant parasites portrays the developmental bifurcation and reveals a regulatory cascade of putative gene functions in determination and subsequent differentiation of each sex. A male-determining gene with a LOTUS/OST-HTH domain points towards unexpected conservation of molecular mechanisms of gametogenesis in animals and a distantly related eukaryotic parasite.

## Introduction

The transmission of malaria parasites to their mosquito vectors requires that a subset of bloodstage parasites switch from repeated asexual replication to sexual development. Epigenetically controlled expression of the transcription factor AP2-G is essential for asexual parasites to commit to sexual development (*1–4*), but the events that regulate the subsequent differentiation of sexually committed parasites into either sex are not understood. Although induced overexpression of AP2-G is sufficient to reprogram asexual parasites for sexual development experimentally (*5, 6*), it remains unknown how this single transcription factor generates two diverging gene expression programs that rapidly lead to male and female gametocytes.

Gametocyte sex ratios differ both between infected hosts and during the course of individual infections in ways that affect transmission to the vector and thereby the epidemiology of malaria (*7–10*). Gametocyte sex determination cannot involve sex chromosomes or inherited mating type loci, because asexual *Plasmodium* blood stages are haploid, and because competent clones retain the ability to produce both male and female gametocytes. Together, these observations suggest an epigenetic, i.e. non-chromosomal, mechanism for the emergence and differentiation of different sexes that is responsive to environmental regulation. Although sex is thought to have evolved only once in the ancestral eukaryote (*11*), mechanisms for how different sexes are determined evolve rapidly (*12*). As a result, none of the genes involved in equivalent determinations in other eukaryotes (*12*) have clear homologues in the Apicomplexa, the phylum of divergent eukaryotes to which malaria parasites belong. The genes involved in sex determination in *Plasmodium* are therefore likely to require employment of a genome-wide screen to be discovered.

In the rodent parasite *P. berghei*, pools of barcoded mutants can now be screened to discover gene functions in an unbiased manner (*13–15*), and blood-stage data suggest that phenotypes observed in *P. berghei* are largely predictive of those in *P. falciparum* (*13, 16*). Here, we have used this approach to screen for genes required for the formation of male and female gametocytes. We then used bulk and single-cell transcriptomics to map the differentiation of these lineages at high temporal resolution. We find that parasites express markers indicative of their eventual sex early in the developmental bifurcation and, by disrupting these genes and characterising mutants, we identify essential components of the male and female transcriptional programs.

## Results

### A systematic screen identifies sexual development genes

To screen for genes required for sexual development, we mutagenized the *P. berghei* reporter line 820 (*17*), which expresses green and red fluorescent proteins (GFP, RFP) from promoters specific for male and female gametocytes, respectively (Fig. 1A). This line was transfected with pools of barcoded *Plasmo*GEM vectors (*18*) targeting 1,302 genes previously determined to be non-essential for asexual erythrocytic growth (relative growth rate at least half of wild type according to (*13*)). Using expression of fluorescent proteins as proxies for sexual development, parasitised red blood cells from mice infected with each superpool were sorted into male (GFP-positive), female (RFP-positive) and asexual (Hoechst-only) populations (~10^6^ of each; Fig. 1B). Barcodes were then counted by deep sequencing of amplicons obtained from the genomic DNA of the sorted parasites. After six replicate screens in large pools, 50 top hits were re-screened in duplicate and with a similar number of control mutants. This smaller pool provided more precise measurements of reporter expression (see Table S1 for all screen data).

**Fig. 1.**
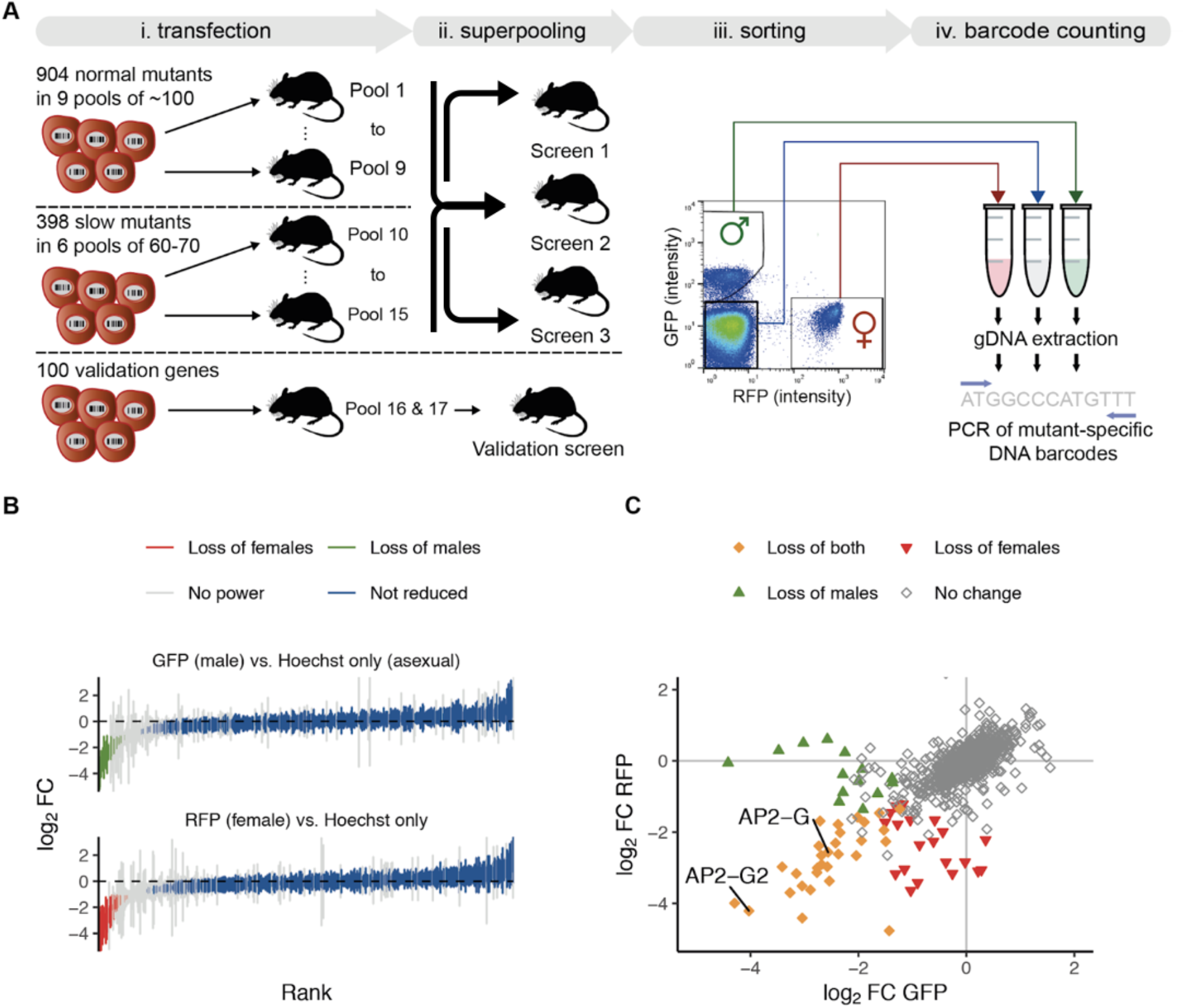
A barseq screen for sexual reporter expression in *P. berghei*. **(A)** Schematic overview showing how *Plasmo*GEM vectors were allocated to pools for transfection and the resulting mutants combined into superpools for sorting on reporter expression. Duplicate barcode PCRs were performed for each sorted population and converted into sequencing libraries for barcode counting. **(B)** All robustly quantified mutants are ranked by the degree to which cells expressing either the male or the female reporter gene were underrepresented. Error bars show standard deviations from at least 4 independent screening experiments. **(C)** Combined results from both reporters, showing filled symbols where the underrepresentation was significant for either one or both sexes. Two highlighted mutants in ApiAP2 genes confirm the expected loss of both markers from the population, as published (*4*).

96% of parasite genes that could be queried were not required for the expression of sexual reporter proteins because their respective barcodes were equally represented in the sorted populations (Fig. 1B). At the defined significance threshold, 30 mutants were depleted from both sexual populations, another 14 were reduced only in the male and 21 only from the female population (Fig. 1C, Table S1). Reassuringly, hits included the transcriptional activator *ap2-g* (*4, 6*)and the repressor *ap2-g2* (*4, 19*), both known to regulate gene expression during gametocyte formation in *P. berghei*. Other genes affecting both sexes often had weaker effects and were also required for normal asexual growth ((*13*); Table S1), suggesting they contribute to cell survival more broadly, examples including a putative NAD synthase (PBANKA_0827500) and a pre-mRNA splicing factor (PBANKA_0409100). Only a few mutants resembled *ap2-g* in affecting both sexual markers profoundly, while not reducing asexual growth. Most notable in this category were a putative ubiquitin-conjugating enzyme (PBANKA_0806000) and a conserved *Plasmodium* protein of unknown function (PBANKA_0824300). The notion that both sexes require these genes for fertility is consistent with evidence from an earlier screen showing that neither the female nor the male gametocyte can pass either of these disrupted alleles to the oocyst stage, which establishes the infection in the mosquito (>24-fold reduction in oocyst numbers shown by (*15*); Table S1).

Genes with functions specific to a single sex were highly represented among a small group of 60 genes that we previously showed respond within six hours of experimentally reprogramming ring stages to sexual development by induced expression of AP2-G (*6*). Since early response genes may hold a clue to sex determination, we wanted to increase the temporal resolution of the time course experiment. *In vitro* synchronised schizonts were reprogrammed into gametocytes, injected into mice and harvested for bulk RNA-seq analysis at additional time points during the first 6 hours after induction (Fig. S1, Table S2). Co-expression analysis by neural network-based dimensionality reduction now identified an even smaller cluster of 12 coregulated genes that responded to *ap2-g* induction within 1-2 h and plateaued from 8-12 h (Fig. 2, cluster 32, and Table S2). This group contained no previously known sex markers, and their transcripts increased before the main wave of around 300 canonical male and female specific genes, whose expression only began to increase detectably from 8-12 h after induction (Fig. 2A). Single sex screen hits accounted for five of the early response genes, a significant enrichment (p < 10^-8^). Cluster 32 included several putative nucleic acid binding proteins of unknown function (Fig. 2B), which we hypothesized could be involved either in determining the sex of gametocytes or in their subsequent sex-specific differentiation. To examine this idea further, we selected all five screen hits from this cluster for further validation. We added to the validation group (shown in Fig. 2C), three genes from other clusters, which also responded rapidly to *ap2-g* overexpression. Cluster 32 included two additional genes encoding putative nucleic acid binding proteins, PBANKA_1302700 and PBANKA_1454800, which had, however, not been covered by the screen because they lacked barcoded *Plasmo*GEM vectors. These genes were selected to complement the hits from the unbiased screen because their domain architecture and expression pattern suggested they may be functionally related. Genes in the validation set are not required by asexual blood stages according to available growth data (*13*).

**Fig. 2.**
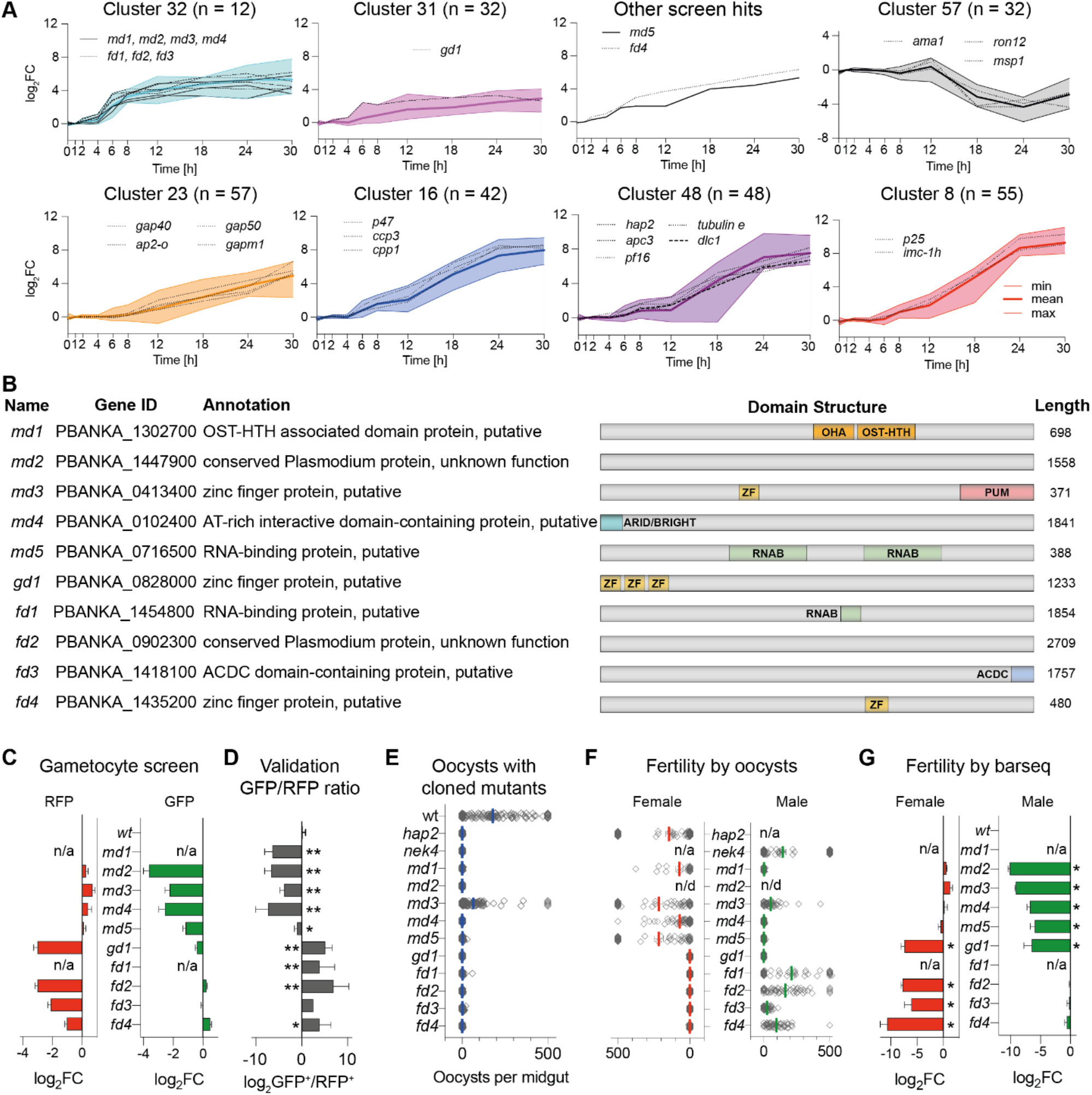
Selection and validation of ten *P. berghei* genes with sex-specific roles in gametocytogenesis. **(A)** Selected gene expression clusters from a bulk RNA-seq timecourse of induced sexual development. Ring stage parasites were reprogrammed at t = 0 h by inducing *ap2-g* (*6*). Relative transcript abundances are given as log2-fold change relative to uninduced, asexually developing parasites. Shown are selected clusters [*number of genes*] with screen hits designated *md, fd* or *gd*. Selected well-characterised marker genes of male, female and asexual development are also shown. **(B)** Schematic illustration of genes with validated roles in sexual development. OST-HTH = oskar-TDRD5/TDRD7 winged helix-turn-helix domain; OHA = OST-HTH associated domain; ARID/BRIGHT = AT-rich interaction domain; ZN = C3H1 zinc finger; PUM = Pumilio RNA-binding repeat profile; RNAB = RNA-binding domain; ACDC = apetala 2 domain-coincident C-terminal domain; PH-like = PH domain like. **(C)** Change in reporter-positive cells in the barseq screen. Error bars show standard deviations. p < 10^-10^ for the affected sex. **(D)** Sex ratio in individual mutants determined by flow cytometry. Error bars show standard deviations from 2-4 biological replicates with cloned mutants, except for *fd3*, where the uncloned population is shown. * p < 0.05; ** p < 0.01 in unpaired T test. **(E)** Transmission efficiency of mutant clones in vivo determined by counting oocysts on midguts 10 days after an infectious blood meal. **(F)** Male and female fertility as determined by the ability of mutant clones to give rise to oocysts in mosquitoes when crossed to *nek4* and *hap2* mutants, which provide fertile male or female gametes, respectively. Oocysts counts show combined data from 25-80 dissected mosquitoes from 2-3 independent experiments. n/d = not done. n/a = not applicable. **(G)** Change in female and male fertility determined by barseq of infected midguts following mutagenesis of female-only or male-only lines, respectively. Error bars show standard deviations from four biological replicates. * p < 0.001.

Flow cytometry with individual knockouts in the 820 line confirmed the biased expression of fluorescent sex reporters for all screen hits and further showed sex-specific losses of marker expression for the two newly included early response genes (Fig. 2D, Fig. S2). Depending on the affected sex, we refer to the validated genes as “male development” (*md1* to *md5*) or “female development” (*fd1* to *fd4*, Fig. 2B). As expected, mutants showed a complete or nearly complete loss in their ability to form oocysts in mosquitoes, with the exception of *md3*, in which oocysts were merely reduced (Fig. 2E). Genetic crosses using either individual mutants (Fig. 2F) or barcoded single-sex pools (Fig. 2G) showed that for most genes, fertility was sex-specifically affected precisely as predicted by reporter expression. Two notable deviations from the screen results were observed. A cloned mutant in PBANKA_0828000, which by FACS only lacked parasites expressing the female marker, additionally suffered from male infertility (Fig. 2F); due to its broader gametocyte development (gd) phenotype we refer to this gene as *gd1*. The second mutant requiring further consideration is *md3*, for which both a transmission experiment (Fig. 2E) and a cross with a cloned line (Fig. 2F) showed that fertile male gametocytes were still present. However, consistent with the screen result, male fertility of all male mutants, including *md3*, was reduced to <1% when measured in competition with fertile mutants (Fig. 2G). We hypothesise that the *md3* mutant produces fertile microgametes at a level that is much reduced, but still sufficient to give rise to a relative abundance of oocysts, probably because optimised laboratory infections of *P. berghei* produce an excess of zygotes, so that oocyst numbers begin to saturate at low input levels (*20*).

### Males and females differentiate from a shared sexual branch

To characterize the developmental block in each mutant more precisely, we performed a series of single-cell RNA sequencing (scRNA-seq) experiments. Using the plate-based Smart-seq2 method, we generated single cell transcriptomes from 2028 red blood cells infected with mutant parasites, and 689 wild-type controls for comparison. These parasites had all been cultured for 24 h to allow any atypically developing gametocytes to survive without being cleared by the spleen (Tables S1 and S3, Fig. S3A and B). We combined this Smart-seq2 data with a high-resolution map of gametocyte development created from 6191 droplet-based single cell transcriptomes covering the asexual cycle, sexual commitment, and the bifurcation into either sex. The two datasets were integrated to generate a combined uniform manifold approximation and projection (UMAP) plot (Fig. 3A and S4). Branching and pseudotime analysis on the combined data showed male and female gametocytes initially follow a common transcriptional trajectory after branching from the asexual cycle, before they assume distinct sexual identities (Fig. 3A, S5 and S6).

**Fig. 3.**
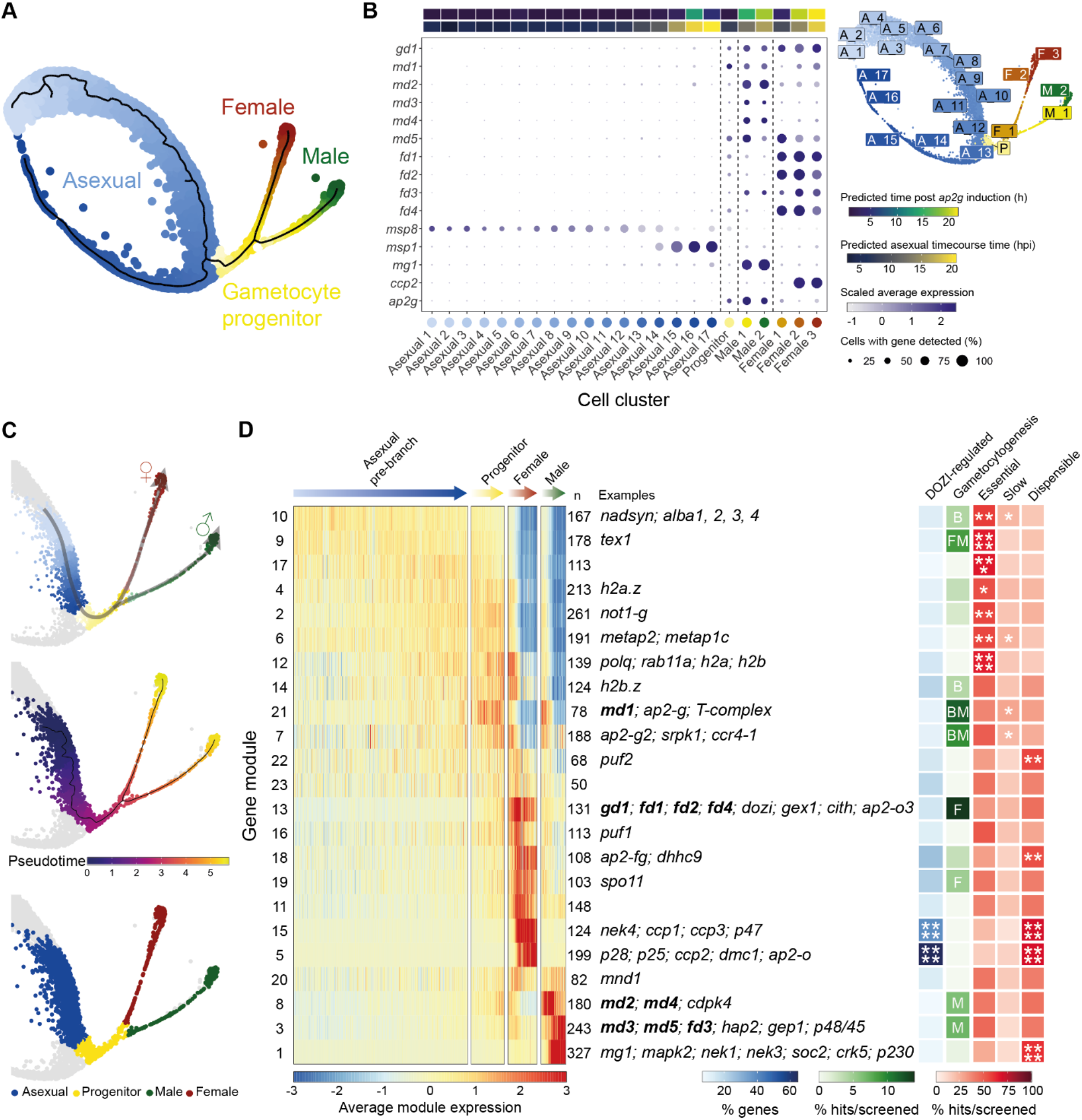
Combined analysis of wild-type 10x and Smart-seq2 transcriptomes from *P. berghei* infected mouse red blood cells. **(A)** UMAP plot of all 6880 wild-type cells (Smart-seq2 and 10x), coloured by their assigned sex designation and progression along development. **(B)** Dot plot showing expression of known marker genes alongside the candidate genes in 23 cell clusters. The estimated average time point of each cluster is annotated by correlating single-cell data to bulk time course data (this study and (*21*)). Cluster locations are shown on the UMAP plot (top right). **(C)** Wild-type cells were sub-clustered and 10x-only cells were selected in order to re-calculate pseudotime for the branches of interest and construct modules of genes that are co-expressed over pseudotime. **(D)** Heatmap showing the scaled average expression of gene modules in cells shown in C. n = the number of genes per module. DOZI-regulated = % of DOZI-regulated genes within each cluster according to (*22*). B, F, and M, represent significant (P ≤ 0.05) enrichment of both, female, or male hits in the screen, respectively. Essential, Slow, Dispensable = % genes per cluster with asexual blood stage phenotype according to (*13*). Examples include previously characterised *Plasmodium* genes with: female-specific expression (*ap2-o3* (*23*), *ap2-fg* (*24*), *nek4* (*25, 26*), *dhhc9* (*27*), *ccp2* (*25*), *ccp1* (*28, 29*), *ccp3* (*29, 30*), *p25* (*31-33*), *p28* (*32*)); functions in meiosis (*spo11* (*34*), *spo11-2* (*34*), *dmc1* (*35*), *gex1* (*36*)); male-specific expression (*cdpk4* (*37*), *hap2* (*38*), *mg1* (*26*), *soc2* (*39*), *crk5* (*39*)); female-biased RNA-binding proteins (*puf1* (*40*), *puf2* (*41*), *dozi* (*42*), *cith* (*17*)); All 8 T-complex proteins, except *cct4* (*30*). * = P ≤ 0.05; ** = P ≤ 0.01; *** = P ≤ 0.001; **** P ≤ 0.0001.

We clustered the 10x transcriptomes to resolve the branch points of sexual development (Fig. 3B). Transcripts from most sexual development genes first became detectable at the joint root of both sexes (Fig. 3C, Table S5 & Fig. S6B, bipotential cluster), where *ap2-g* was also upregulated, but before transcripts of canonical sex genes such as a dynein heavy chain (*mg1*, male) and *ccp2* (female) became detectable. *md* and *fd* genes were generally upregulated most strongly along the specific sexual trajectory affected by their disruption and thus serve as early markers downstream of *ap2-g* (Fig. 3B). Having assigned cells to male, female and asexual lineages in pseudotime (Fig. 3C), we identified co-expression gene modules in the single-cell data that delineate the sex-specific developmental programs (Fig. 3D). Mapping all knockout phenotypes from the screen onto these gene modules shows a significant enrichment for gametocytogenesis phenotypes among the first wave of sex specific genes in each branch (p < 10^-3^), (Fig. 3D, Table S4), providing further validation for the screen and independent confirmation of the early response genes first identified in the bulk transcriptomes from the reprogramming time course (Fig. 2A and S1).

### Single cell transcriptomes distinguish differentiation genes from putative regulators of sex ratio determination

Using the wild-type data to stage parasites (Fig. S7), we found that mutants fell into two classes: those where cells of the infertile sex were undetectable and those where the infertile sex was still present but had an atypical transcriptional signature (Fig. 4A, B). The latter category of mutants still expressed some of the core marker genes of the infertile sex but cells often clustered separately from wild type (Fig. S8, S9 and Table S6), and more cells resembled earlier points in pseudotime (Fig. 4B). This analysis identified *md4, md5, fd2, fd3* and *fd4* as differentiation mutants, because they become committed to a sex but fail to develop its complete transcriptional signature. Whilst the loss of *gd1* results in near complete absence of females, it also perturbs differentiation in males (Fig. 4A and B). Differential gene expression analysis shows differentiation mutants to be defined by unique transcriptional states in the infertile sex (Fig. 4C, S9 and Table S6), suggesting each gene affects transcription through a unique mechanism.

**Fig. 4.**
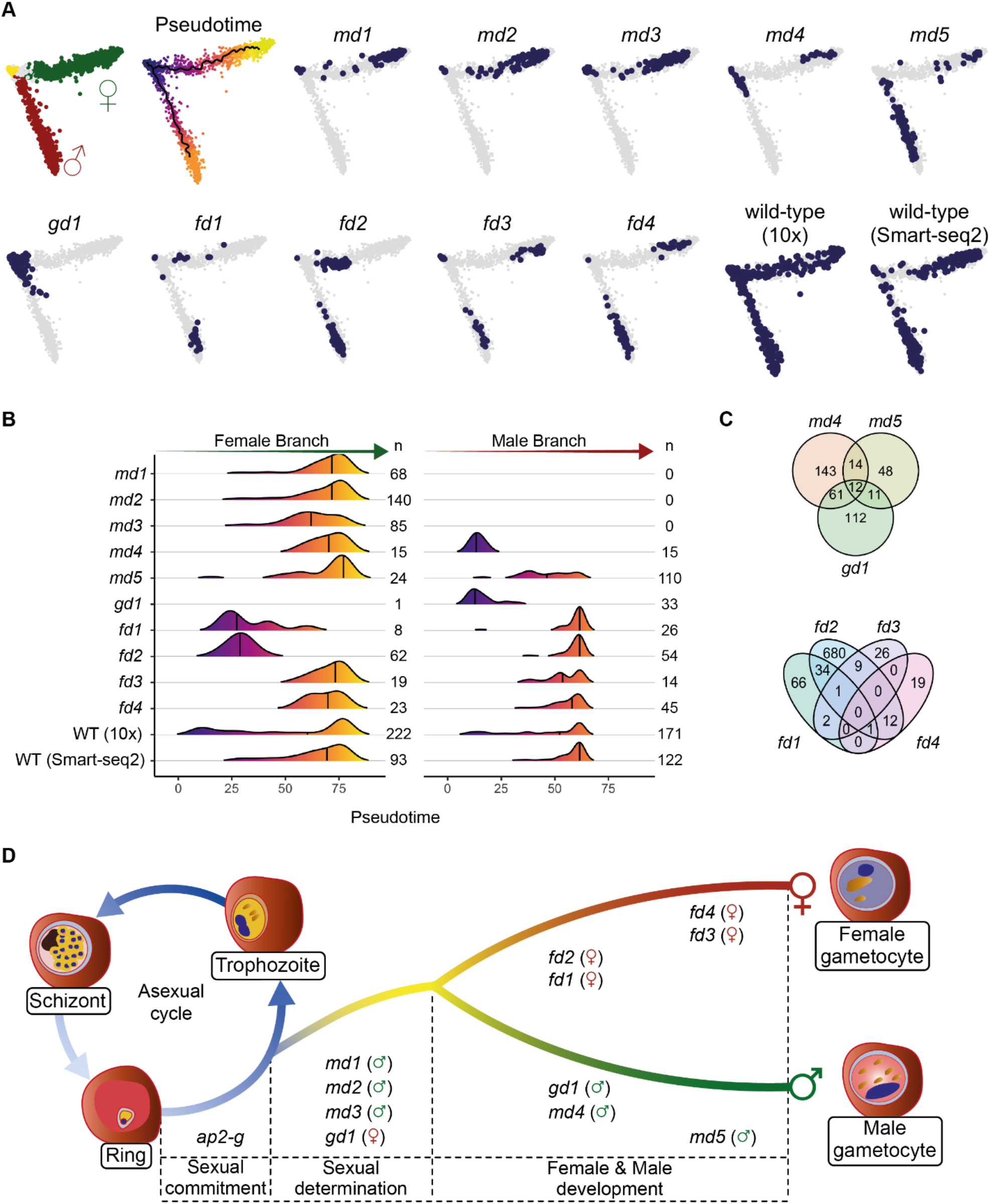
Smart-seq2 analysis of *in vitro*-matured wild-type and mutant parasites. **(A)** Principal Component Analysis (PCA) plots of 3012 transcriptomes from single parasitised RBCs of the sexual branch that were obtained by merging all wild-type and mutant cells and subsetting the branch of interest (Fig. S7) **(B)** Density plots showing the distribution of assigned female (left) and male (right) cells along each pseudotime trajectory and grouped by genotype. n = the number of cells for each condition as shown in panel A. WT = wild-type. The line indicates the median. **(C)** Venn diagrams showing the numbers of genes with differences in transcript abundance in female and male gametocytes, respectively, relative to wild-type. Genes that result in the absence of a sex cannot be evaluated. **(D)** Model of how sexual determination and differentiation is affected by a cascade dominated by putative nucleic acid binding proteins identified in this study. In addition to its role in commitment, *ap2g* probably has sex specific roles in differentiation (*90*), together with sex-specific transcription factors, such as *ap2-fg* (*24*).

When the sex ratios of mutants were re-assessed using Smart-seq2 data for staging (Fig. 4B), four candidate genes for sex ratio determination emerged. *md1* and *md2* are potential maleness-determining genes since their disruption leads to a complete loss of cells expressing male markers. *md3* also showed a marked sex ratio shift towards females. However, consistent with the reduced male fertility retained by this mutant (Fig. 2E and F), we found enough *md3* males for enrichment by flow sorting. The transcriptomes of these cells were essentially indistinguishable from wild type (Fig. S10), suggesting deletion of *md3* does not affect male differentiation, but sex ratio, albeit with less complete penetrance than *md1* and *md2*. *gd1* behaved like a sex determining gene with respect to the female pathway, because cells with a female transcriptional signature were absent; the inability of a disrupted *gd1* locus to transmit to oocysts (Fig. 2E) may thus involve different mechanisms in each sex.

*md1* is the most noticeable AP2-G responsive gene to be upregulated at the base of the sexual branch (Fig. 3C, D). It encodes a protein with a putative LOTUS/OST-HTH domain (*43, 44*). OST-HTH domain proteins exist in pro- and eukaryotes, but only those of animals have been studied in detail and all are involved in gametogenesis (*45*). Examples include Oskar, an important germline determining factor in *Drosophila* embryos (*46*) and the tudor domain contain proteins (TDRD5/TDRD7) with sex-specific fertility functions in mice (*47*). LOTUS domains organise ribonucleoprotein (RNP) complexes and it may therefore be significant that the zinc finger gene *md3* contains a weak homology with a pumilio RNA-binding domain, while *md2* lacks clear homologs outside of *Plasmodium*. Together, these considerations raise the intriguing possibility that creating the male lineage in *P. berghei* shares elements with germline definition in multicellular organisms.

Male differentiation mutants are *gd1, md4* and *md5* in order of decreasing severity with respect to the presence of core male transcripts (Fig. S9). Disrupting *gd1* and *md4* has profound but distinct effects on male gene expression. *md4* encodes a conserved *Plasmodium* protein of unknown function characterized by a putative N-terminal ARID/BRIGHT DNA binding domain, which in other eukaryotes targets developmental transcription factors to AT-rich DNA sequences (*48*), suggesting it may regulate transcription downstream of the initial commitment to the male developmental trajectory. Transcript abundance of *md4* decreases in mature males but it stays high in *gd1* mutants (Fig. S11), illustrating the early developmental arrest of *gd1* males and also indicating that *gd1* is not required for the expression of *md4*. Deletion of *md5* has a less severe effect on core male transcripts and likely operates through a different mechanism because the protein contains putative RNA binding motifs.

Among the female differentiation genes, *fd1* encodes a putative RNA-binding protein, and *fd2* encodes a conserved *Plasmodium* protein. Both have profoundly perturbed female transcriptomes, and *fd1* mutant females expressed only some female markers. In contrast, *fd3* and *fd4* have more subtle roles for the formation of transcriptionally normal females. *fd3* is characterized by an AP2-coincident C-terminal domain of unknown function (ACDC). This type of domain was first identified in a number of *P. falciparum* AP2 domain-containing proteins (*49*), but in *fd3* it is not associated with a known DNA-binding domain. *fd4* encodes another putative zinc finger protein, and the mutant only shows a moderate downregulation of a few late female transcripts (Fig. 4C, S11 and Table S6). This gene is downregulated in *fd1* and *fd2* mutants, suggesting it may operate downstream of both.

## Discussion

Our transcriptomic data suggest a model in which male and female gametocytes differentiate from a common sexual precursor in ways that rely on a cascade of nucleic acid binding proteins which are co-expressed downstream of AP2-G and fulfil distinct functions in a hierarchy of regulatory events (Fig. 4D). In this hierarchy the triple zinc finger protein GD1 is the strongest candidate for a top-level factor for female determination or differentiation upstream of the female-specific transcription factor AP2-FG (*24*). *Fd1 to fd4* probably contribute downstream and with less impact (Fig. 4D).

Additional work is required to identify how sex ratio is determined in *P. berghei*. One view of how the sexes are formed in *P. falciparum* is that commitment to a particular sex co-incides with or even precedes the induction of sexual development by AP2-G (*50*). In that case, gene expression in response to AP2-G would be expected to follow a sex-specific pattern from the start. Importantly, neither our global transcriptomic analysis of single cells, nor the expression of candidate genes from the functional screen produced evidence that a sex-specific transcriptomic signature precedes the commitment to sexual development. Instead, we observed a common branch of sexual precursors and discovered key roles for AP2-G early response genes, suggesting sex may be determined downstream of *ap2-g* induction. Alteratively or additionally, non-transcriptional mechanisms may operate upstream of *md1-3* and *gd1*, involving for instance chromatin marks, differential splicing or phosphorylation states that determine the emergence of sex-specific transcriptional signatures downstream of AP2-G. Notwithstanding its early role in the switch to sexual development, AP2-G also binds to the upstream sequences of many male and female specific genes later during gametocytogenesis.

Genes that respond early during the induction of sexual development have also been identified in *P. falciparum*. All ten *P. berghei* genes validated here have orthologs in *P. falciparum* (Table S7), where their transcripts are all upregulated during sexual development, with seven transcripts peaking during early (stage I-II) gametocytogenesis (*51*). Furthermore, chromatin immunoprecipitation of AP2-G in synchronously developing parasites identified *P. falciparum* orthologs of *gd1, fd1, fd2, md3*, and *md4* as likely direct targets for AP2-G binding in stage I gametocytes, but not in sexually committed ring stages or asexual schizonts (*52*). Taken together these data point at a high degree of conservation in the first steps of gametocyte differentiation following the initial commitment to sexual development in malaria parasites.

In summary, our functional screen, in combination with single-cell transcriptomes of cloned mutants, has identified a diverse group of proteins that are co-expressed downstream of AP2-G and whose deletion affects either the determination of, or differentiation along, a male or female cellular trajectory. Further analysis of these genes will shed light on the precise molecular mechanisms of sex ratio determination and sexual development. The screen revealed an abundance of putative RNA binding proteins, including the maleness inducing factor MD1 and the early response genes MD4, MD5 and FD1, whose molecular targets now need to be identified. The LOTUS/OST-HTH domain gene *md1* raises the intriguing possibility that the function of this domain in gametogenesis is conserved beyond animals, possibly to the origin of sex in the ancestral eukaryote. Sex determination mechanisms evolve rapidly but often involve RNA-dependent regulation, for instance through differential splicing or translational repression (*53–55*) and our data suggest similar principles operate in *P. berghei*. Discovering the molecular targets and interactors of the proteins of GD1, MD1 and FD1 now provide a route to establishing these mechanisms in more detail.

## Acknowledgements

The authors would like to thank the staff of the Illumina Bespoke Sequencing and Core Cytometry teams at the Wellcome Sanger Institute for their contribution.

## Funding

Work at the Wellcome Sanger Institute was funded by Wellcome core grant 206194/Z/17/Z awarded to OB, M.L. and A.J.C.R. Work at Umeå University received funding from the Knut and Alice Wallenberg Foundation and the European Research Council (Grant agreement No. 788516). Work at the University of Glasgow (APW, ABR, RSK) was funded by the Wellcome Trust (Refs: 083811, 104111 & 107046 to APW and the BBSRC (Ref BB/J013854/1 to RSK). RSK is supported by BBSRC (Ref BB/J013854/1). KKM is supported by Wellcome Trust and Royal Society (202600/Z/16/Z). MH is supported by SNF (P2SKP3_187635) and HFSP (LT000131/2020-L).

## Competing Interests

The authors declare they have no competing interests.

## Data and materials availability

The raw scRNA-seq data for this study have been deposited in the European Nucleotide Archive (ENA) at EMBL-EBI under accession number PRJEB44892) (https://www.ebi.ac.uk/ena/browser/view/PRJEB44892) and raw bulk RNA-seq data are available from the NCBI Gene Expression Omnibus (GEO) (accession number GSE110201 and GSE168817). Supporting files and code are available on Github at https://github.com/andyrussell/Gametocytogenesis. Screen data can be visualised on the *Plasmo*GEM website https://plasmogem.shinyapps.io/Gametocytes_Shiny/. Single-cell RNA-seq data can be searched and visualised on the Malaria Cell Atlas website www.malariacellatlas.org. Mutant scRNA-seq data can be interacted with on the cellxgene instance http://obilab.molbiol.umu.se/gcsko.

## Supplemental information

Materials and Methods

Supplementary Tables S1 - S8

Supplementary Figures S1 - S14

## Notes

https://www.ebi.ac.uk/ena/browser/view/PRJEB44892

https://plasmogem.shinyapps.io/Gametocytes_Shiny/

https://www.malariacellatlas.org

http://obilab.molbiol.umu.se/gcsko

### Competing Interest Statement

The authors have declared no competing interest.

