## Supplemental Material for "Regulators of male and female sexual development critical for transmission of a malaria parasite"

**Supplementary materials**

**Materials and Methods**

**Gene targeting vectors and primers**

All gene targeting vectors used in the barseq screen were obtained from the *Plasmo*GEM resource (http://plasmogem.sanger.ac.uk/search) (*18*), from where details of all vector designs and sequences of gene-specific primers are also available. The barseq screening vectors carried the default *Plasmo*GEM 3xHA-hdhfr-yfcu gene replacement cassette (*56*), and are itemised in Table S8. The *Plasmo*GEM vectors used to generate single gene KOplasmogem lines are also listed in Table S8.

Since *Plasmo*GEM vectors were not available to target *md1 and fd1*, knock-out vectors were prepared using PCR. For md1 amplicons of 1.25 kb upstream and downstream of the gene were amplified from genomic DNA and then assembled either side of a selection cassette amplified from the *Plasmo*GEM 3xHA-hdhfr-yfcu gateway vector (*56*) by Gibson Assembly using primer overhangs. The Gibson product was used as the input for a PCR reaction to amplify the entire construct, which was then gel-purified prior to transfection.

For *fd1* a CRISPR/Cas9 knock-out vector was constructed by PCR amplification of 0.5 kb 5' and 3' regions of the coding sequence of the target gene from genomic DNA, which were restriction-ligation cloned so to flank an *eef1α* 5'UTR-*tgdhfr-CAM* 3'UTR resistance cassette in a vector also holding the U6 RNA Polymerase 3 promoter from *Plasmodium yoelii* to drive expression of the target specific guide RNA (gRNA) cloned in by BsmBI, to generate plasmid ABR063 (Fig. S12). The fd1 CRISPR/Cas9 knock-out line was generated in a split-Cas9 line where conditional activation of CAS9 is achieved by rapamycin-induced dimerisation of a C-terminal fragment of Cas9 fused to FK506- and rapamycin-Binding Protein domain (C-Cas9-FKBP) and a N-terminal fragment of Cas9 fused to the FKBP-Rapamycin Binding domain (N-Cas9-FRB). To achieve this line, N-Cas9-FRB and C-Cas9-FKBP was cloned so to flank the bidirectional 5'UTR of eef1α, and to become nested within 5' and 3' targeting sites for the p230p locus generating plasmid ABR010 (Fig. S12). Primer sequences for cloning custom knock-out vectors and genotyping the resulting parasites are displayed in Table S8.

For scRNA sequencing experiments, where possible, FACS sortable knockout vectors were generated by converting *Plasmo*GEM intermediate vectors (*14*) into knockout vector with a gateway cassette containing an pbhsp70 5'utr-GFPmut3-pbdhfr 3'utr-hdhfr-yfcu expression cassette for constitutive expression of GFP upon vector integration, which allows for FACS sorting to select wild-type free ko parasites without the need for dilution cloning. The gateway construct was generated by amplifying a 1.4 kb fragment of the 5'utr of hsp70 from *P. berghei* genomic DNA , which was cloned into the R6K-GFPmut3-hdhfr-yfcu Gateway vector (added to the *Plasmo*GEM tagging vector repertoire described in (*14*), upstream of GFPmut3 to generate the R6K-hsp70p-GFPmut3-hdhfr-yfcu Gateway vector. When no FACS-sortable knockout vector could be generated in this way, scRNA-seq analysis was performed on dilution cloned mutants generated using the standard *Plasmo*GEM knockout vectors harbouring the 3xHA-hdhfr-yfcu cassette (*md2* and *md3*) or custom knockout vectors (*md1* and *fd1*).

**Parasite lines**

The mutants assayed in the the barseq screen and phenotyped by FACS, in mosquito feeds or scRNA-seq experiments as dilution cloned single gene knockout lines were generated in the *P. berghei* 820cl1m1cl1 line (referred to as 820) that expresses GFP under the control of a male-gametocyte-specific promoter and RFP under the control of a female-gametocyte-specific promoter (*17*, *26*). For the scRNA-seq experiments that utilised the FACS sortable knockout vectors, the knockout lines were generated in the non-fluorescent *P. berghei* ANKA cl15cy1 line (*57*). These GFP positive knockout lines were selected for by FACS and assayed together with $mCherry_{hsp70}$ wild-type parasites that constitutively express mCherry, also under control of the hsp70 5'utr but from within a modified p230p locus and are described in (*58*). For scRNA-seq of the target genes that lacked fluorescence sortable knockout vectors, the 820 knockout lines were assayed after dilution cloning and without the internal wildtype control since fluorescence could not be used to infer knockout versus wild type genotype by sorting. *P. berghei* 820 knock-out lines were dilution cloned by standard limited dilution and all single gene knock-out parasite lines were genotyped by diagnostic PCR on gDNA as described in (*13*) (Fig. S13).

The *fd1* knockout line was made using a new *P. berghei* 820 split-Cas9 line (G1262Cl1), generated by transfecting construct ABR010 into a 820 Gene In Marker Out (GIMO) ready parasite line G879Cl1 (*6*). In this new 820 split-Cas9 line, G1262Cl1, the P1 site of p230p harbours the 820 expression cassette with male specific GFP expression and female specific RFP expression and the P2 site of p230p contains the split-Cas9 cassette described above.

**Use of rodents**

All animal research at the Wellcome Sanger Institute was conducted under licenses from the UK Home Office, and protocols were approved by the Animal Welfare and Ethical Review Body of the Wellcome Sanger Institute. Rats were housed as two cage companions and mice as five cage companions. They were housed in individually ventilated cages (IVC) furnished with autoclaved aspen woodchip, fun tunnel and Nestlets at 21 ± 2°C under a 12:12 hr light dark cycle at a relative humidity of 55 ± 10%. Rodents were kept in specific-pathogen-free conditions and subjected to regular pathogen monitoring by sentinel screening.

Female RCC Han Wistar outbred rats (Envigo, UK) aged seven to sixteen weeks were infected with *P. berghei* parasites by intraperitoneal injection. Infected rats served as donors for *ex vivo* schizont cultures typically on day four to five of infection, at a parasitemia of ~1%–5%. Rats were terminally anaesthetized by vaporized isoflurane administered by inhalation prior to terminal bleed. Rats were used because they give rise to more schizonts with higher transfection efficiency compared to mice. Transfection efficiency is critical when screening pools of vectors.

Mice used at the Wellcome Sanger Institute were bred in-house or purchased from Envigo, UK. To generate barseq pools and superpools transfected parasites were injected intravenously into the tail of female adult BALB/c inbred mice aged six to eleven weeks. There is one exception, which is that some pools of slow mutants were grown in seven week old female SCID ($Prkdc^{scid}$) mice (Table S9). The Balb/c animal model was chosen to minimize host genetic variability and to obtain robust infections with a low incidence of cerebral pathology. SCID mice were selected due to their lack of adaptive immunity to improve the representation of slow mutants that are commonly lost during passage. Generation of single gene ko lines, revival of frozen stabilates and dilution cloning was carried out in either adult female BALB/c mice, or adult female outbred TO (HsdOla:TO) mice, which also generate robust infection with low rates of cerebral malaria. FACS analysis and scRNA-seq analysis of single gene ko lines was performed in female BALB/c aged eight to fourteen weeks. Genetic crossing experiments and mosquito feeds were performed using female TO mice and backbites using female C57BL/6N mice. C57BL/6N mice were used for backbites as they are the most susceptible to malaria infection by sporozoites.

The animal research at Umeå University was conducted under Ethics Permit A13-2019 and approved by Swedish Board of Agriculture (Jordbruksverket). Mice were group-housed as four cage companions in IVC with autoclaved woodchip, paper towels for nesting and a paper fun tunnel or kidney dish, at 21 ± 1°C under a 12:12 h light dark cycle at a relative humidity of 55% ± 5%. Specific-pathogen-free conditions are maintained and subjected to Exhaust Air Dust (EAD) monitoring and analysis biannually. The mice used at Umeå University were purchased from Charles River Europe. Female Balb/c aged 6-20 weeks old were used to bring up stabliates, carry out dilution cloning, do mosquito feeds and to perform FACS analysis of single gene ko lines.

All animal work in Glasgow was approved by the University's Animal Welfare and Ethical Review Body and by the UK's Home Office (PPL 60/4443). The animal care and use protocol complied with the UK Animals (Scientific Procedures) Act 1986 as amended in 2012 and with European Directive 2010/63/EU on the Protection of Animals Used for Scientific Purposes. Mice in Glasgow facilities are held in groups of up to 5 per cage, in IVC, containing standard woodchip/aspen, sizzle nesting, fun tunnels and dome homes. Room temp is 21.5 ± 1°C with a 12:12 light dark cycle and a relative humidity of 50% ± 6%. Specific-pathogen-free conditions are maintained and subjected to analysis annually. Parasites were maintained in Theiler's original (TO) or NIH Swiss outbred female mice, approximately weighing 25 g and > 6 weeks old.

Animals at all sites were fed a commercially prepared autoclaved dry rodent diet and water, both available *ad libitum*. The health of animals was monitored by routine daily visual health checks. The parasitemia of infected animals was determined by methanol-fixed and Giemsa-stained thin blood smears.

#### Generation of mutant superpool

To allow genome-scale phenotyping in a single experiment by assaying of all mutants in a single mouse we created superpools of all viable *P. berghei* mutants using pools of vectors matched for integration efficiency and asexual fitness in (*13*). All genes identified as dispensable in the asexual screen were allocated into 9 groups based on their normalised abundance on day 6 of the infection, which was taken to represent vector integration efficiency. Genes giving slow asexual growth in the asexual screen were allocated into three groups based on their relative growth rate, and then each of these groups was further divided into two based on normalised abundance on day 6. This gave a total of 9 pools of dispensable genes (Pool 1-9: 100-110 vectors per pool) and 6 pools of genes generating slow-growing mutants (Pool 10-15: 65-70 vectors per pool), (Table S8). Vectors were picked for all pools and prepared in a pooled midi-prep approach as discussed previously (*13*).

All pools were transfected individually into rat-derived 820 *P. berghei* schizonts and injected intravenously (IV) into mice. Transfectants were selected by 0.07 mg/mL pyrimethamine administered in drinking water, all as previously described (*13*, *59*). At a parasitaemia of 1-5% (day six to seven post-trans infection for dispensable mutant pools 1-9, and day seven to nine for slow mutant pools 10-15) infected blood was collected and frozen down.

**Screening of mutant superpool**

Stabilates of mutant pools were thawed and combined into different superpools of normal and slow growing parasites such that each vector was part of four independent screening experiments. Fig. 1A and Table S9), and then immediately injected IV into mice and propagated under pyrimethamine selection. On day 6-7 of the infection (parasitaemia ~10%), a sample of (~100 µL) infected blood was collected as "input" and the rest of the blood was collected directly into 4 mL gametocyte non-activation medium (RPMI1640 with L-glutamine, without phenol red and sodium bicarbonate (Sigma) supplemented with 0.1% BSA, 4mM sodium bicarbonate and 20 mM Hepes at a pH of 7.25) at room temperature. White blood cells were removed by passing through Plasmodipur filters (Proxima) and gametocytes were subsequently purified on a Histodenz (Sigma) density gradient (13.25% w/v). Following purification, parasites were washed and stained with Hoechst 33342 (Thermo Fisher) prior to sorting. Throughout handling, care was taken to avoid gametocyte activation by not exposing infected blood or parasite pellets to air, performing all steps in the gametocyte non-activation medium and keeping all reagents at room temperature.

GFP+ Hoechst+ (male gametocytes), RFP+ Hoechst+ (female gametocytes) and Hoechst+ only (asexual parasites) populations were isolated and sorted using a BD Influx cell sorter. Typically 0.5-1.0 $x10^6$ cells were collected for the GFP+ Hoechst+ and RFP+ Hoechst+ populations and 1.0-2.0 $x10^6$ cells were collected for the Hoechst+ populations from each sort. Following sorting, cells were collected by spinning at 2000 x *g* for 10 min, in 5 mL Eppendorf tubes, supernatant was carefully removed and pellets frozen at -20 °C. For the input infected blood sample, erythrocytes were lysed using $NH_4Cl$ and parasites were pelleted by centrifugation; the supernatant was removed and pellets were frozen at -20°C. Genomic DNA was extracted from the pellets upon thawing using phenol-chloroform. gDNA from sorted gametocyte populations was reconstituted in 10 µL $dH_2O$. For input infected blood sample, gDNA was dissolved in 100 µL. 5 µL gDNA was used as input for the first PCR reaction when generating sequencing libraries, with each reaction run in duplicate. Barcode sequencing libraries were generated using a nested, direct-amplification PCR approach with Illumina index tag primers to allow multiplexing of all samples from one experiment. Libraries were sequenced on a MiSeq (Illumina) at cluster density of 400 K with 50% PhiX spiked-in, all as previously described (*14*).

**Targeted screen**

The screen was adapted to re-screen all gametocyte phenotype hits from the initial genome-scale screen (forming the "targeted screen"). This targeted screen pool contained <166 mutants (Table S8) and its mutant pools were generated by transfecting two pools of 83 constructs into two separate mice as described as above. Resulting stabilates were mixed together and re-injected into mice (Table S9). For the targeted screen conducted *in vivo*, the experiment was performed exactly as outlined above. Gametocytes were purified and sorted with mutant barcodes sequenced as before.

**Screen analysis**

Barcodes in Illumina sequencing data were counted and tabulated per library. To facilitate downstream analysis, these counts were expressed as $\log_2$ proportions, with 0.5 added to counts. To quantify the uncertainty induced by the fact that the barcodes that successfully entered libraries were a random sample of those in the actual population (with sampling occurring at the time of purification, sorting, pipetting of template material, and during amplification), standard deviations were calculated for each barcode in each sample, based on technical PCR duplicates. These standard deviations were made more precise by taking the moving average of values (k=11), with samples ordered by the abundance of the barcode in question, and enforcing monotonicity. This approach provided an estimate of the $\log_2$ proportion made up by each barcode in each sample, with an associated uncertainty estimate. For each pool analysed, results were available based on sorted populations derived from three mice. We calculated a fold-enrichment for each sample of interest (the fluorescently sorted populations) as compared to the negative control (the Nycodenz gradient for super pool 3 (SP3), and specifically sorted non-fluorescent parasites for super pool 4 (SP4) & the targeted screen super pool 6 (SP6)), and propagated uncertainties to this value. To normalise these values within each sample, we calculated an estimate for the change in abundance of seven control genes (PBANKA_071830, PBANKA_120780, PBANKA_083110, PBANKA_051060, PBANKA_051820 & PBANKA_132610), and computed an inverse-variance weighted mean for the change in this control sample (all inverse-variance weighted mean calculations used the method described in (*13*)). We then normalised by subtracting this control value from each value of interest. To yield a combined estimate across all mice and experiments, we calculated the inverse-variance weighted mean, and its expected uncertainty using the method described in (*13*)**.** Named phenotypes were assigned using thresholds based on the upper and lower bounds of the confidence intervals for enrichment in the GFP-positive and RFP-positive populations: if the confidence interval overlapped a two-fold reduction in barcode counts, we considered that we did not have power to detect a substantial reduction (*no power*), if the confidence interval lay entirely with a reduction of more than two-fold, we considered this a significant reduction (*reduced*) and if the confidence interval lay entirely with a reduction of less than two fold, we considered this to represent no substantial reduction.

**Single knockout transfections and genotyping**
Single gene ko *Plasmo*GEM vectors for md2, md3, md4, md5, gd1, fd2, fd3 and fd4 were prepared using QIAGEN Plasmid Midi Kit and 1-5 ug NotI digested and ethanol precipitation purified DNA was transfected into *P. berghei* 820 rat-derived schizonts. Transfected parasites were injected intravenously into mice and selected using pyrimethamine for pooled transfections above. For *md1* and *fd1*, 1 µg of custom vector DNA was transfected into schizonts as above, with the single modification that mouse-derived schizonts were used.

**FACS of single gene ko lines**
FACS was performed either on parasites revived from frozen stabilates (typically on day 5 post-injection at a parasitaemia of >2 %) or direct from transfection (typically on day 7-8 post-transfection at a parasitaemia of >2 %). Gametocytes were analysed directly from infected mouse blood or were purified from using gametocyte non-activation medium and Histodenz density gradient, and stained with Hoechst 33342 as above. Purified and stained gametocytes were resuspended in gametocyte non-activation medium, taking care to not expose parasite pellets to air and kept at room temperature at all times. Samples were immediately analysed for

mCherry (female gametocytes) and GFP (males gametocytes) using a BD LSR Fortessa instrument and data were analysed using Flowjo (v. 7.6.5 and 10.6.1) .

**Mosquito infections and genetic crosses**

Stablilates with *P. berghei* 820 dilution cloned single ko lines (md1, md2, md3, md4, md5, gd1, fd1, fd2, fd3 and fd4) or wt parasites were revived and then infected blood was collected by cardiac puncture. For single parasite lines mice were directly infected by intraperitoneal injection. For genetic crosses *md1, md3, md4, md5, gd1, fd1, fd2, fd3* and *fd4* knockout lines were mixed at a 1:1 ratio of iRBC with a *nek4* knockout (*60*) or a *hap2* knockout (*38*) prior to infection with the mixed parasite infected blood intraperitoneally. On day three post-infection (typically at 2-10% iRBC) parasitaemia was assessed by microscopic observations of Giemsa stained blood films and exflagellation evaluated as previously described (*38*). Infected mice were anesthetised and ~50 mosquitoes allowed feed on each mouse for 15-20 min at 19°C. Unfed mosquitoes were removed after 24 hours. Midguts were dissected and oocysts were counted using light microscopy on day nine to twelve days post-feeding. Backbites were performed only for those single ko feeds that produced oocysts

**Bulk-RNA-seq**

The generation and initial analysis of the additional time points for the bulk RNA-seq dataset was performed as described previously (*6*). Briefly, the $PB_{GAMi}$ line (engineered to overexpress the AP2-G transcription factor and undergo synchronous conversion into gametocytes upon the induction with rapamycin) was synchronised to the late schizont stage and either induced or treated with vehicle only. At different time points post induction, the blood containing the developing parasites was harvested. Plasmodipur filters (EuroProxima) were used to remove the leukocytes according to the manufacturer’s instructions and red blood cells were lysed by resuspension in ice-cold 1× E-lysis buffer (1.5M $NH_4Cl$, 0.1M $KHCO_3$, 0.01 EDTA). The resulting parasite pellet was washed with 1xPBS and stored in Trizol reagent for future RNA extraction. Independently, in order to generate the reference transcriptome of male and female gametocytes, the 820 reporter line (*17*), was used to sort $5x10^6$ male and female gametocytes, as described in the previous section. The resulting male and female pellets were also stored in Trizol for further processing. Complete RNA was isolated from all the samples using Trizol/chloroform extraction followed by isopropanol precipitation and 1-2 µg of starting material was taken for mRNA isolation and stranded RNA-seq library construction. The libraries were prepared using NEBNext library preparation modules, with minor modifications of the protocol shown to improve the yield when sequencing AT-rich transcriptome of *Plasmodium* parasites (*61*). The samples were pooled and sequenced using an Illumina HiSeq 2500 system according to the manufacturer’s instructions. All samples were generated in biological duplicates or triplicates and uninduced controls were always generated and processed together with the induced samples.

**Smart-seq2 scRNA-seq**

*Cell preparation and staining*

Knockout mutants for *md4*, *md5*, *gd1*, *fd2*, *fd3*, and *fd4* were created in a background constitutively expressing GFP. Each mutant was individually mixed in a 1:1 ratio (based on parasitemia counts) with $mCherry_{hsp70}$ wild-type parasites constitutively expressing mCherry as internal control. Inclusion of a co-infecting wild-type strains allows to control forenvironmental influences on sex ratio, sexual commitment, as well as transcriptional variation between hosts. Mutants created in the *P. berghei* 820 background (*md1*, *md2*, *md3*, and *fd1*) cannot be combined with an internal mCherry control, but an external mCherry control in another mouse was included (Fig. S3B). All mice were treated the same from this point onwards. 3 days after inoculation, mice were terminally bled by cardiac puncture. The ~1 mL blood sample was immediately transferred into a pre-warmed (37 °C) 1.5-mL tube, and transferred into a sealed culture flask containing 50 mL of schizont culture medium (RPMI with 20% FBS, 15 mM $NaHCO_3$, and 1% penicillin/streptomycin). Parasites were cultured for 24 hours at 36.5 °C with shaking at 65 rpm. Cultures were harvested by centrifugation at 450 x *g* for 3 minutes at room temperature. Late-stage- and gametocyte- infected red blood cells were purified on a 55% Histodenz gradient by centrifugation at 300 × *g* for 20 minutes at room temperature. Cells were washed once and resuspended in 1 mL gametocyte non-activation media.

*Cell sorting*

Sorting was performed as described in (*62*). Briefly, 4 uL of lysis buffer (0.8% of RNAse-free Triton-X (Fisher) in nuclease-free water (Ambion)), UV-treated for 30 min with a Stratalinker UV Crosslinker 2400 at 200, 000 µJ/cm2, 2.5 mM dNTPs (Life Technologies), 2.5 µM of oligo(dT) (Non-anchored OligoDT, HPLC purified, 100 µM, 5'AAGCAGTGGTATCAACGCAGA GTACTTTTTTTTTTTTTTTTTTTTTTTTTTTTTT3'; IDT) and 2U of SuperRNAsin (Life Technologies)) was dispensed in 96-well plates. Cell sorting was performed on an Influx Cell Sorter (BD Biosciences) with a 70 µm nozzle. For mixed parasite populations,cells were sorted by gating for single-cell events, Hoechst positive events (compared to an uninfected RBC control), and then on GFP (mutant population) or mCherry (wild-type population). For 820-background parasites, this strategy was modified so that parasite cells were sorted by gating for single-cell events and on GFP (male), mCherry (female), or Hoechst (unbiased sort of all parasites of that genotype). A non-sorted negative control, and a positive 100-cell control were included on every plate.

*Library preparation and sequencing*

First and second strand cDNA synthesis and pre-amplification were performed as described in (*62*) with 96-well plates and 25 PCR cycles. Quality control of cDNA samples was monitored using a high-sensitivity DNA chip on the Agilent 2100 Bioanalyzer. Libraries were prepared using dual indexes as described in (*62*) and pools of 384 cells (4 plates) were sequenced onto one lane on a HiSeq 4000 using v4 chemistry with 75 bases paired-end reads and run according to manufacturer's instructions.

**10x Genomics Chromium scRNA-seq**

$mCherry_{hsp70}$ parasites were used for all experiments, so any data generated was comparable with Smart-seq2 data which used this background as a wild type control. Blood was obtained as previously described (3 days post infection), a leukodepletion step was included by using a pre-wetted Plasmodipur syringe filter (EuroProxima) was used for leukodepletion prior to culturing. To cover the whole span of sexual development and mitigate unequal representation due to sequestration, two separate cultures from two mice were set-up in a staggered way so that they could be harvested at the same time, respectively 30 minutes and 12 hours after blood harvest (Figure S2C).

Cultures were smeared prior to harvesting in order to ascertain precise parasitemia. After harvesting, the total number of red blood cells in each sample was counted using a single-use hemocytometer (NanoEntek). This count was corroborated using a Countess cell counter. The concentration of infected red blood cells (iRBCs), derived from the parasitaemias and red blood cell concentration, in each culture was established and cells were pooled 1:1. Cells were loaded according to the manufacturer's instructions to recover 5000 cells. Chromium 10x v2 chemistry was used and the library was prepared according to manufacturer's instructions and sequenced across 2 lanes of a HiSeq 2500 on Rapid Run settings using asymmetric paired-end sequencing (26 cycles for Read 1 and 98 cycles for Read 2)..

**Bulk RNA-seq data analysis**

The raw data processing, generation of initial *.cram files and adapter removal was performed using the default analysis pipelines of the Sanger Institute. The raw data was transformed into paired *. fastq files using Samtools software (ver. 1.3.1) (*63*). The generated reads were re-aligned to the *Plasmodium berghei* ANKA genome (PlasmoDB-30 release) in a splice-aware manner with HISAT2 (*64*) using the --known-splicesite-infile option within the splicing sites file generated based on the current genome annotation. Resulting *.bam files were sorted and indexed using Samtools (ver. 1.3.1) and HT-seq python library (ver. 1.3.1) (*65*). was used to generate reads counts for all genes for further processing.

The matrix of gene counts was combined with the time points published previously (*6*). Differential expression analysis was performed at each time point between the induced and uninduced samples using R (v3.4.4) with DESeq2 package v 1.18 (*66*). In parallel, the differential expression analysis was performed between male and female gametocytes as well as between each gametocyte sex and asexual parasites. In order to cluster the genes according to their responses to the AP2-G induction and sex specificity, the fold differences in gene expression at each time point as well as fold differences between male and female gametocytes were extracted from DEseq2 expression tables and used as input for the self-organising maps training algorithm implemented in the “kohonen” R package (ver. 2.0.14) (*67*). The ‘som’ algorithm was run with 8x8 map size, 200 data presentation cycles and default “alpha” parameters. The relative expression of each of the 64 clusters was visualised using plotting functions implemented within the “kohonen” package. In order to confirm the separation of male and female expression clusters, genes classified as female-, male-, gametocyte- and asexual-specific, based on the differential expression were overlaid on the map in in order to visualise the clusters they belonged to. Genes were classified as male/female/asexual specific if they were differentially upregulated eg. females (with log2FC >2 and FDR<0.05) when compared to both asexual parasites and males. Genes overexpressed in gametocytes but without clear preference between the sexes were classified as gametocyte-specific. Full list of genes with their cluster assignments in Table S2.

The significance of the cluster based on the sex-specific genes was calculated using the hypergeometric probability density function (P),

$$P(\text{k,N,K,n}) = \frac{\binom{K}{k}\binom{N-K}{n-k}}{\binom{N}{n}} \quad (1)$$

$$pvalue = \sum_{k}^{K} P(x) \quad (2)$$

where *k* and K are the numbers of sex-specific genes and the number of total genes in a given cluster, respectively, and n and N are the total number of sex-specific genes in the screen and the total number of genes in screen, respectively. For each cluster the p-value was calculated by sum over probabilities for greater or equal than the enriched number of sex-specific genes to test the null hypothesis (equation 2).

**Mapping and generation of expression matrices for scRNA-seq data**

*Smart-seq2 mapping*

Single-cell *Plasmodium* transcriptomes were mapped as reported previously (*62*). CRAM files were downloaded from iRODs Wellcome Sanger Institute core pipeline. CRAM files were converted to FASTQ format using Biobambam2 (v2.0.37) (*68*) (*bamtofastq exclude=SECONDARY,SUPPLEMENTARY,QCFAIL*). Nextera adaptor sequences were trimmed using Trim Galore (v0.4.3) (*69*) (*trim_galore -q 20 -a CTGTCTCTTATACACATCT --paired --stringency 3 --length 50 -e 0.1*). Trimmed FASTQ files were then mapped using HISAT2 (v2.1.0) (*70*) and indexes were produced using the *P. berghei* v3 genome sequences (*71*), downloaded from GeneDB (*72*) (October 2016). Trimmed reads were then mapped using default parameters (*hisat2 --max-intronlen 5000 -p 12 -q -x*). GFF files were downloaded from GeneDB (October 2016) and converted to GTF files using an in-house script. All feature types (mRNA, rRNA, tRNA, snRNA, snoRNA, pseudogenic_transcript and ncRNA) were conserved, with their individual 'coding' regions labelled as CDS in every case for convenience. Where multiple transcripts were annotated for an individual gene, only the primary transcript was considered. Reads were summed against genes using HTSeq (v0.11.2) (*65*) *(htseq-count -f bam -r pos -s no -t CDS*). HTSeq excludes multimapping reads by default (-a 10). This means that reads mapping ambiguously to similar genes from the same family are not considered in our analysis.

*10x data alignment, cell barcode assignment, and UMI counting*

The sequencing reads in CRAM format were downloaded from iRODs Wellcome Sanger Institute core pipeline. CRAM files were converted to FASTQ format using the samtools fastq command (*63*). Cell Ranger (version 2.1.1) was used to create a reference file from the *P. berghei* v3 genome (obtained from: www.sanger.ac.uk/resources/downloads/protozoa/) *(71)* using standard parameters (*cellranger mkref --genome=.. --fasta=.. --genes=..*). The gene PBANKA_0713500 was manually corrected to include two exons with missing ids. FASTQ files were passed into the Cell Ranger 2.1.1 workflow to assign each read to a cell barcode and UMI using standard parameters (*73*) (*cellranger count --id=.. --transcriptome=.. -fastq=..*).

**Filtering and normalization of scRNA-seq data**

R version 4.0.3 (2020-10-10) was used for all scRNA-seq analysis (*74*).

*Smart-seq2 filtering*

Count matrices and associated metadata (phenodata) were read into R (version 4.0.3) and no cell and 100 cell controls were removed from further analysis. Seurat (version 3.2.2) was used for pre-processing (*75*). 63/5245 genes were not detected in any cell and were also removed. Cells with genes per cell < 220, genes per cell > 3300, percentage of total counts mapping to mitochondrial genes > 20%, and number of counts per cell < 1000 were removed (Fig. S14A). This resulted in the removal of 706/3450 cells.

*10x filtering*

The raw output barcodes, genes and matrix files were read into Seurat (v3.2.2) using the Read10X command (*75*). To distinguish cells from background, an expectation–maximization (EM) algorithm was applied using mixtools (v.1.2.0) (*76*) to discover where the two distributions (cells and background) intersected. This resulted in the identification of 7762 cells (Fig. S14B). Low-quality cells were then filtered by removing any cell that contained <200 genes per cell; this removed 1131 cells resulting in 6631 cells (Fig. S14C).

*Normalization and doublet detection*

In both datasets cells were normalized (*NormalizeData(x, normalization.method = "LogNormalize", scale.factor = 10000*), variable genes were found (*FindVariableFeatures(x, selection.method = "vst", nfeatures = 2000*), and data was scaled (all.genes = all genes in the datset; i.e. *ScaleData(x, features = all.genes)*). For the 10x data, doublets (cell barcodes that are associated with a significant number of reads from multiple cells) were filtered out using DoubletFinder v3 (*77*) (*doubletFinder_v3(pb_sex, PCs = 1:21, pN = 0.25, pK = 0.01, nExp = nExp_poi, reuse.pANN = FALSE, sct = FALSE*). Doublet removal was not applied to the Smart-seq2 dataset as a singlet gate was applied during FACS.

**Single-cell transcriptome analysis of wild-type data**

*Data integration*

Wild-type cells from the Smart-seq2 and 10x datasets were integrated using Seurat v3.2.2 (*75*). Each dataset was subsetted to only include genes that were present in both (5018 shared genes). Datasets were then individually normalised (*NormalizeData()*) and variable features were found (*FindVariableFeatures(x, selection.method = "vst", nfeatures = 2000*). Anchors were found (*FindIntegrationAnchors(object.list = x, dims = 1:21*) and integration was performed (*IntegrateData(anchorset = x, dims = 1:21, features.to.integrate = shared.genes; where shared.genes is all 5018 genes*).

*Cell projection, clustering, and annotation*

To identify subpopulations of cells in the integrated dataset, PCA was performed (RunPCA(x, npcs = 30) and then cells were projected into two dimensions using the UMAP algorithm (*78*) using the first 10 principal components after inspection of an elbow plot to detect significant principal components in Seurat v3.2.2; the other parameters used were: n.neighbors = 150, min.dist = 0.4, repulsion.strength = 0.03, local.connectivity = 150. Cells were clustered in a ten dimension UMAP space using the Louvain algorithm with multilevel refinement at a resolution of 2 (*79*).

Cells were manually annotated by assigning each Louvain cluster to either asexual, progenitor, male or female, based on established marker genes (Fig. S5). All wild-type cells were then ordered along pseudotime using Monocle 3 (version 0.2.3.0) (*80–82*) (learn_graph(x, learn_graph_control=list(ncenter=550, minimal_branch_len = 15), use_partition = FALSE; order_cells(x)). The root cells were selected manually using the interactive feature. The median pseudotime value for each cluster was then used to order clusters within each lineage. At this point, two female clusters collapsed into one due to their similar median pseudotime.

The progenitor cluster was analysed further (n = 206, all cells were from the 10x dataset). Variable features were calculated for this subset and the cells were projected into 30-dimension PCA space. The elbow plot of these PCs was inspected and seven of these dimensions were used to find neighbours. Clusters were generated using the Louvain algorithm with multilevel refinement at a resolution of 0.5 (*75*) (Fig. S6A). Marker genes for these resulting two clusters were found using the MAST framework implemented in Seurat v.3.2.2 (FindAllMarkers(x, only.pos = FALSE, min.pct = 0.25, logfc.threshold = 0.25, test.use = "MAST") (*83*)

*Pseudotime and gene module generation*

In order to analyse the sexual branch, the following clusters were subsetted for further analysis: Asexual_10, Asexual_11, Asexual_12, Progenitor, Male_1, Male_2, Female_1, Female_2, Female_3. 9 outlier cells were removed from analysis as they were located far from the clusters and would have affected pseudotime calculation. This resulted in a subset of 2817 branch cells (Fig. 3C).

For the pseudotime analysis of the sex branch, the subsetted cells were further subsetted so that only wild-type cells generated using 10x were included in the analysis. The reason 10x cells were only used in following steps was because there is not a robust way to account for the batch effects between the Smart-seq2 and 10x dataset within Monocle and Monocle is not compatible with negative values generated during Seurat batch correction. The subsetted object counts matrix was preprocessed (preprocess_cds(x, num_dim = 50, norm_method = "log")) and UMAP coordinates were extracted from the Seurat object. The graph structure was learnt (learn_graph(x, l learn_graph_control=list(ncenter=550, minimal_branch_len = 30), use_partition = FALSE)). The cells were ordered along pseudotime (order_cells(x)) and the root cells were selected manually using the interactive feature. The 23 gene modules were generated by first performing graph_test(x, neighbor_graph="principal_graph", cores=8, expression_family = "negbinomial"). Significant genes were selected by removing any genes with a *q*-value > 0.05. Gene modules were then found using find_gene_modules(x[significant_genes,], resolution=c(10^seq(-6,2)), random_seed = 1234), to maximise the number of modules as suggested by the developers of Monocle 3.

*Enrichment of gene classes in gene modules*

A list of the gametocyte screen hits was used to find the percentage of hits in each module, which was given by the number of screen hits present in that module divided by the number of genes that were screened in that module. The percentage of DOZI-regulated genes per module was calculated by extracting the genes identified in both of the DOZI and CITH fractions of RIP-Chip experiments performed in (*22*), and calculating the percentage of these genes that appear in each module. The percentage of genes per module for each of the asexual phenotypes identified in (*13*) was also calculated for genes that were covered in that screen. The significance of the module based on the DOZI/CITH-associated genes, gametocyte screen hits, and asexual screen hits, respectively, was calculated using the hypergeometric probability density function (P), as shown above in the Bulk RNA-seq data analysis section. In this case, *k* and K are the numbers of gene hits and the number of total genes in a given module, respectively, and n and N are the total number of hits in the screen and the total number of genes in screen, respectively. For each module the *p*-value was calculated by sum over probabilities for greater or equal than the enriched number of sex-specific genes to test the null hypothesis (equation 2).

**Single-cell transcriptome analysis of single knockout mutant data**

*Data integration*
To compare mutant transcriptomes to wild-type ones, all cells from the Smart-seq2 and 10x datasets were integrated using Seurat v3.2.20 (*75*). Each dataset was subsetted to only include genes that were present in both (5018 shared genes). Datasets were then individually normalised (NormalizeData()) and variable features were found (FindVariableFeatures(x, selection.method = "vst", nfeatures = 2000). Anchors were found (FindIntegrationAnchors(object.list = x, dims = 1:21) and integration was performed (IntegrateData(anchorset = x, dims = 1:21, features.to.integrate = shared.genes; where shared.genes is all 5018 genes).

*Cell projection, clustering, and sex branch isolation*

To identify subpopulations of cells in the integrated dataset, PCA was performed (RunPCA(x, npcs = 30) and then cells were projected into two dimensions using the UMAP algorithm (*78*) using the first 10 principal components after inspection of an elbow plot to detect significant principal components in Seurat v3.20.20; the other parameters used were: n.neighbors = 150, min.dist = 0.4, repulsion.strength = 0.03, local.connectivity = 150. Cells were clustered in a ten dimension UMAP space using the Louvain algorithm with multilevel refinement at a resolution of 0.5 (75) (Fig. S7A).

In order to analyse the sexual branch, 11/26 clusters were subsetted for further analysis (Fig. S7D). 8 outlier cells were removed from analysis as they were located far from the clusters and would have affected pseudotime calculation later. The identities of these 8 cells were checked to ensure that they were not enriched in a specific mutant and they contained a mixture of wild-type and mutant cells. This resulted in a subset of 3012 sex branch cells. The cells were projected into 2 dimensional PCA space and 17 new clusters were generated using the first 11 principal components, and the Louvain algorithm with multilevel refinement at a resolution of 1. Each cluster was then assigned a sex identity based on the position in the PCA plot and the detection of known marker genes. Pseudotime was then calculated using Monocle 3 (version 0.2.3.0) (79–81) by first learning the graph structure (learn_graph(x, learn_graph_control = list(ncenter = 200, minimal_branch_len = 10), use_partition = FALSE)). The cells were ordered along pseudotime (order_cells(x)) and the root cells were selected manually using the interactive feature (Fig. 4). Cells were assigned to one of the three resultant branches using choose_graph_segments() in Monocle 3. Finally, cells were assigned an identity if they belonged to both a cluster that was designated as male, female or progenitor as detailed above and also belonged to the male, female or progenitor branch, respectively.

*Differential gene expression analysis*
In order to assess how mutants impacted gene expression, differential expression analysis was performed using Seurat v3.2.2 and MAST (*83*). In order to compare cells from similar points of development, the latest cluster in development in the sexual branch that the mutant affected with representation of the mutant under comparison was chosen. Within this cluster, the mutant

cells were directly compared to the wild-type cells (Fig. S9). For *fd2,* both clusters 15 and 8 were chosen because the representation of wild-type cells in cluster 15 was too low.

### Supplementary Tables

**Table S1. Screen results**

(see separate file)

---

**Table S2. Bulk RNA-seq**

Transcript abundance in *P. berghei* parasites at 0 - 44 h after initiating infections in mice with *in vitro*-synchronised schizonts. Data are shown as log2-fold change in a population reprogrammed to undergo gametocytogenesis by induction of *ap2-g* (Kent *et al*., 2018), compared with uninduced, asexually developing parasites. All data are from synchronised parasite populations growing in mice. Column N additionally shows the log2 of the male-to-famle ratio of normalised transcript abundance in flow-sorted male and female gametocytes of the Pb820 line.

(see separate file)

---

**Table S3. Related to Fig. 3B and C: Assignment of single cells to clusters and associated meta data.**

(see separate file)

---

**Table S4. Related to Fig. 3D: Assignment of genes to co-expression clusters.** Also show screen phenotypes and phenotype enrichment fdr.

(see separate file)

---

| Gene Name | Phenotype description | Expression in wild-type |
|---|---|---|
| *md1*<br>(PBANKA_1302700) | **Sex ratio – no males.**<br>Mutant sex ratio shifted to female.<br>No cells express core male genes.<br>Females fertile with wild-type transcriptome. | 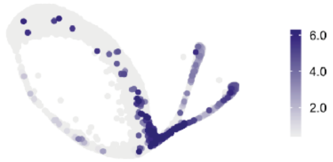 |
| *md2*<br>(PBANKA_1447900) | **Sex ratio – no males.**<br>Mutant sex ratio shifted to female.<br>No cells express core male genes.<br>Females fertile with wild-type transcriptome. | 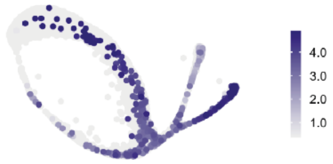 |
| *md3*<br>(PBANKA_0413400) | **Sex ratio – few males.**<br>Mutant sex ratio shifted to female.<br>Only a few males, but these are fertile and have a wild-type transcriptome.<br>Females fertile with wild-type transcriptome. | 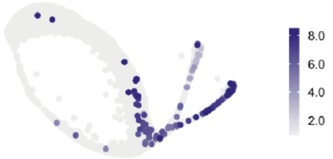 |
| *md4*<br>(PBANKA_0102400) | **Male differentiation.**<br>Infertile males lack transcripts for many core male markers.<br>Perturbed transcriptome distinct from *gd1* males.<br>Other *md* genes expressed. | 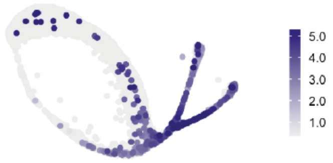 |
| *md5*<br>(PBANKA_0716500) | **Male differentiation.**<br>Infertile males lack transcripts for only some core male markers.<br>Transcriptome distinct from *gd1* and other perturbed males. | 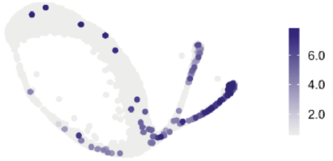 |
| *gd1*<br>(PBANKA_0828000) | **Sex ratio – no females.**<br>Mutant sex ratio shifted to male.<br>No cells expressing core female genes but one borderline cell expressing female early response genes was observed.<br>**Male differentiation.**<br>Infertile males expressing only some core male marker genes. | 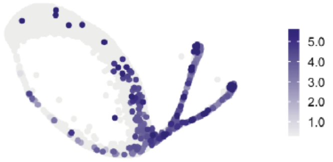 |
| *fd1*<br>(PBANKA_1454800) | **Female differentiation.**<br>Mutant with infertile females lacking transcripts for a unique set of core females markers.<br>*fd2-4* transcripts reduced in mutant. | 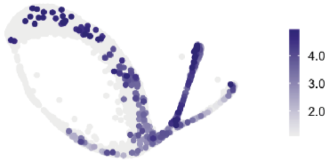 |
| *fd2*<br>(PBANKA_0902300) | **Female differentiation.**<br>Mutant with infertile females lacking transcripts for a different subset of core females markers, overlapping only partly with *fd1*.<br>*fd4* transcript reduced. | 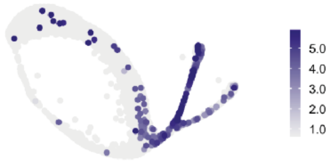 |
| *fd3*<br>(PBANKA_1418100) | **Female differentiation.**<br>Mutant with infertile females but expressing core females markers.<br>Some non-female transcripts increased.<br>Putative role in repression of destabilization of transcripts. | 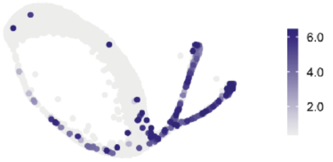 |
| *fd4*<br>(PBANKA_1435200) | **Female differentiation.**<br>Mutant with infertile females expressing most core females markers but lacking some late female transcripts, such as *p28*, *isp1*. | 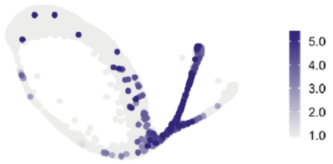 |

**Table S5. Summary data from Smart-seq2 experiments of mutants.** A description of the phenotype observed from the single-cell RNA-seq data is given along with a summary of the fertility phenotype (Figure 2 E, F and G). On the right hand side, scaled expression of the gene in wild-type only cells is shown with the 5th and 95th quantiles set as the minimum and maximum expression values respectively to eliminate any outliers having a strong influence on visualisation.

---

**Table S6.** Smart-seq2 data from wild-type and mutants, including genes with differences in transcript abundance between mutants as determined by Smart-seq2.

(see separate file)

---

| | | | | Direct AP2G target (Josling et al. 2020) | | | van Biljon et al. (2019) |
|---|---|---|---|---|---|---|---|
| ***P. berghei* gene ID** | **Name** | ***Pb* ap2-g early response gene** | ***Pf* 3D7 gene ID** | **stage-I gametocytes** | **sexual rings** | **schizonts** | **Cluster*** |
| PBANKA_1302700 | *md1* | YES | PF3D7_1438800 | no | no | no | 9 |
| PBANKA_1447900 | *md2* | YES | PF3D7_1233200 | no | no | no | 8 |
| PBANKA_0413400 | *md3* | YES | PF3D7_0315600 | YES | no | no | 7 |
| PBANKA_0102400 | *md4* | YES | PF3D7_0603600 | YES | no | no | 8 |
| PBANKA_0716500 | *md5* | no | PF3D7_0414500 | no | no | no | 8 |
| PBANKA_0828000 | *gd1* | YES | PF3D7_0927200 | YES | no | no | 8 |
| PBANKA_1454800 | *fd1* | YES | PF3D7_1241400 | YES | no | no | 8 |
| PBANKA_0902300 | *fd2* | YES | PF3D7_1146800 | YES | no | no | 8 |
| PBANKA_1418100 | *fd3* | YES | PF3D7_1319600 | no | no | no | 8 |
| PBANKA_1435200 | *fd4* | YES | PF3D7_1220000 | no | no | no | 9 |

**Table S7. *Plasmodium falciparum* orthogs of investigated genes and some of their properties.** *P. berghei* early response gene in this study (responding to reprogramming within 1-2 hours). Direct AP2-G target: Has associated consensus binding peak in chromatin immunoprecipitation experiments in *P. falciparum* cultures whose differentiation was induced by AP2-G stabilisation (*52*). Cluster: Co-expression cluster during *P. falciparum* sexual differentiation according to the bulk microarray time course data in (*51*). Cluster descriptions are: 6-10 = increased during gametocytogenesis; 6-7 = transcripts maintained at increased levels throughout commitment and development; 7 = contains many differentiation genes; 8-10 with specific peaks during gametocytogenesis; 8 = transcripts involved in early stage development increasing from stage I-II; 9 = increased from intermediate stage III - IV; 10 = stage V transcripts.

---

**Table S8.**
Vectors and primers
Sheet 1 - Full screen vectors
Sheet 2 - Targeted screen vectors
Sheet 3 - Single gene ko vectors
Sheet 4 - Non-PlasmoGEM primers

(see separate file)

---

**Table S9. Superpooling strategy for the gametocyte barseq screen.**

(see separate file)

### Supplementary Figures

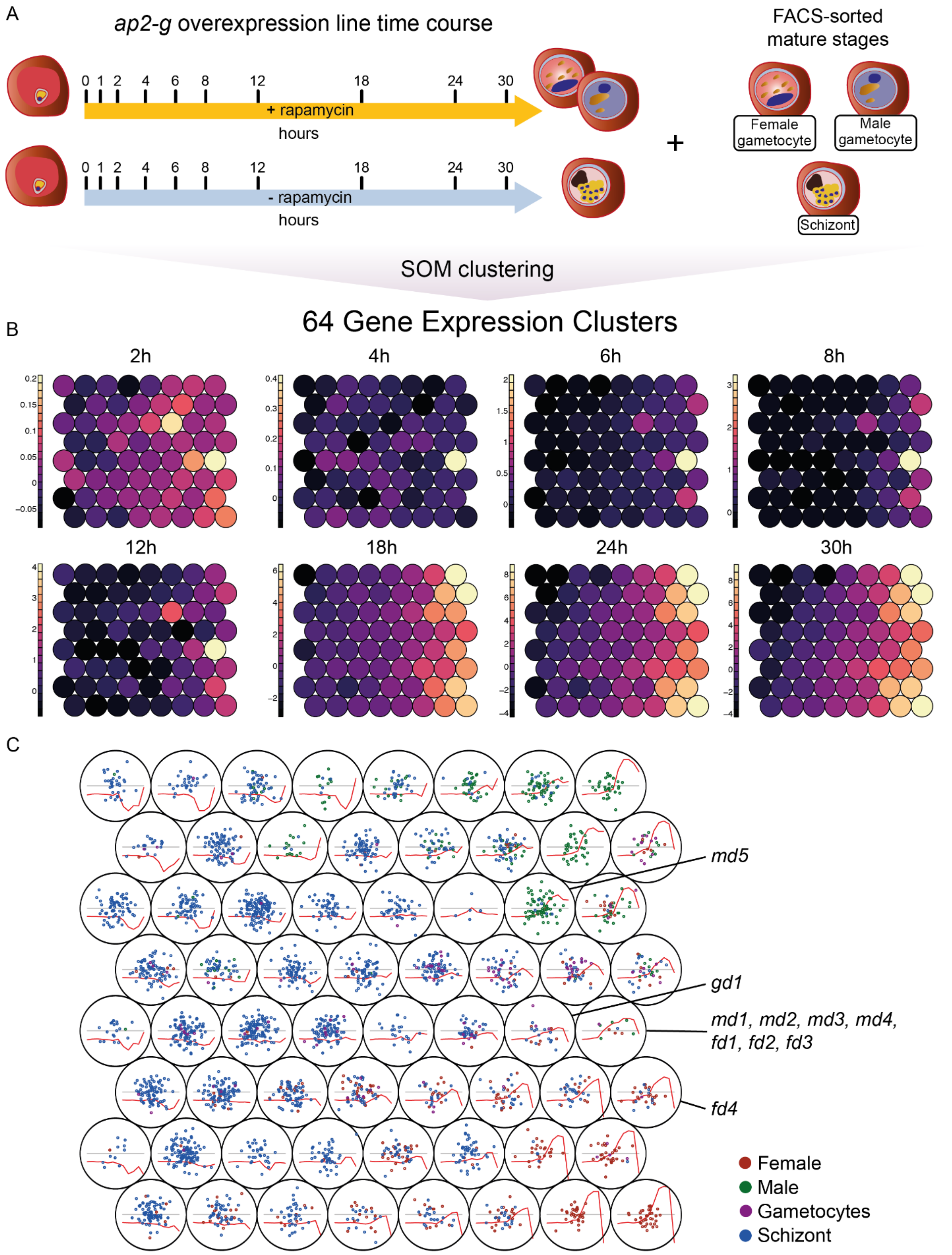


**Fig. S1. Time course of transcriptome changes during induced gametocytogenesis reveals clusters of genes co-regulated during early gametocyte differentiation. (A)** Outline of the experimental design showing samples collected from the induced (+ rapamycin) and uninduced (-rapamycin) ap2-g overexpression line (*6*) as well as purified mature stages controls (schizonts and male/female gametocytes) **(B, C)** Maps generated using the self-organizing map (SOM) algorithm, showing 64 clusters of genes co-regulated during male and female gametocyte development. The relative expression of each of the clusters in the induced/uninduced population is represented by colors **(B)** or red lines **(C)**. Points in **(C)** represent genes classified based on the differential expression between the mature stages as upregulated in schizonts (green), male/female (blue/red) gametocytes or gametocytes of both sexes (purple). Cluster assignment of key sex determinants discussed in the manuscript are marked in **(C)** Cluster assignment and expression levels of individual genes in the Table S2. Lines show the mean relative expression values of the cluster over time.

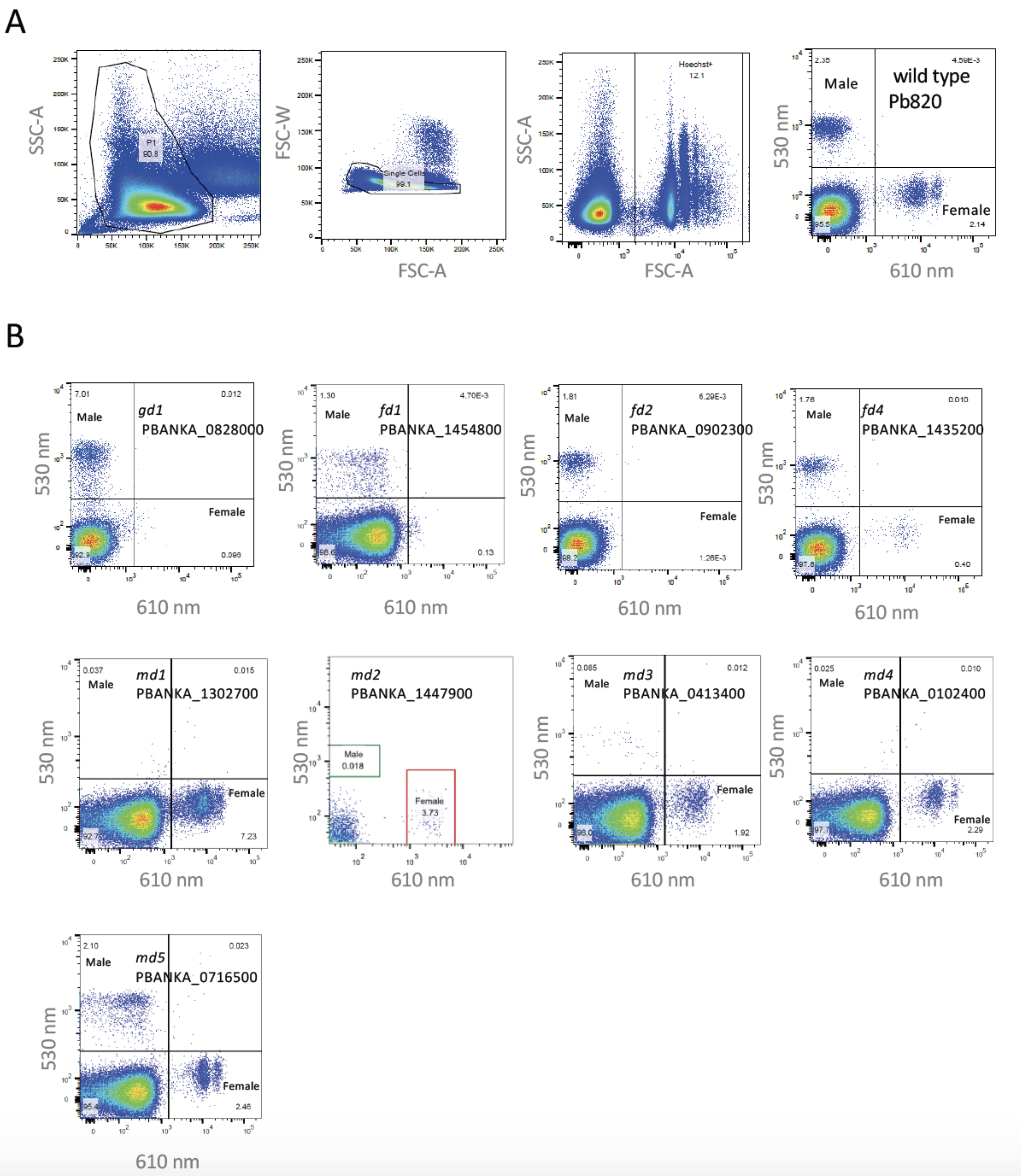


**Fig. S2. Sex ratio analysis of cloned mutants. (A)** Gating strategy for FACS analysis of male and female gametocytes of the Pb820 line and its derivatives: Samples were first gated on Forward Scatter

Area and Side Scatter Area to exclude debris and then gated on Forward Scatter Area and Forward Scatter Width to exclude doublets. Hoechst positive infected red blood cells were then analyzed for mCherry (610 nm, females ) and GFP (530 nm, males ) expression (**B**) Representative FACS plots showing proportion of male gametocytes, female gametocytes and normocytes in the infected red blood cell fraction of different mutants .

---

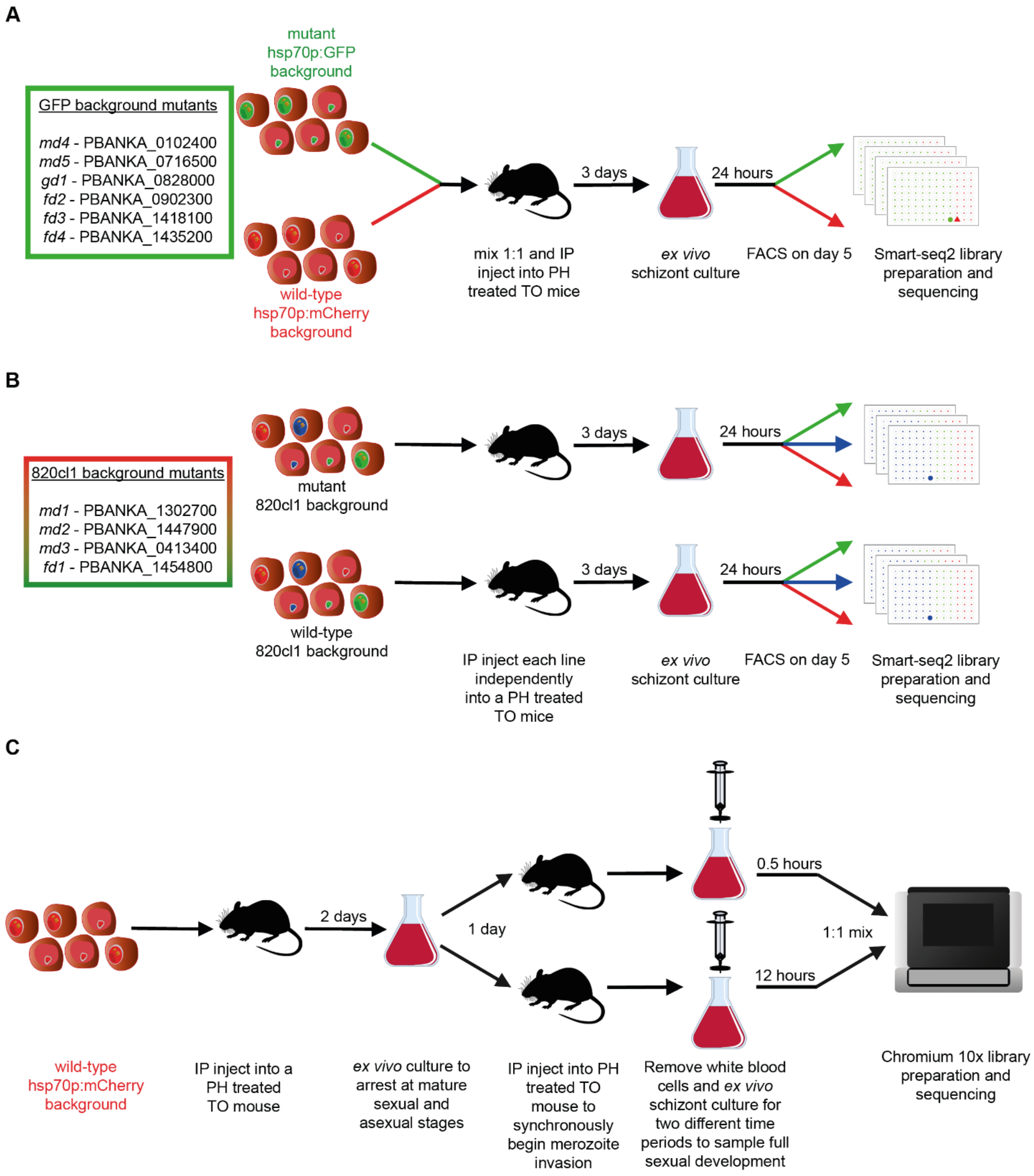


**Fig. S3. Design of scRNA-seq experiments. (A, B)** Smart-seq2 experiments use *in vitro* culture to allow parasites to differentiate along their determined trajectory without risking splenic clearance of developmentally aberrant mutants. Two different backgrounds were used in these experiments. **(B)** 10x experiments were set up to capture cells from intermediate stages of gametocyte differentiation.

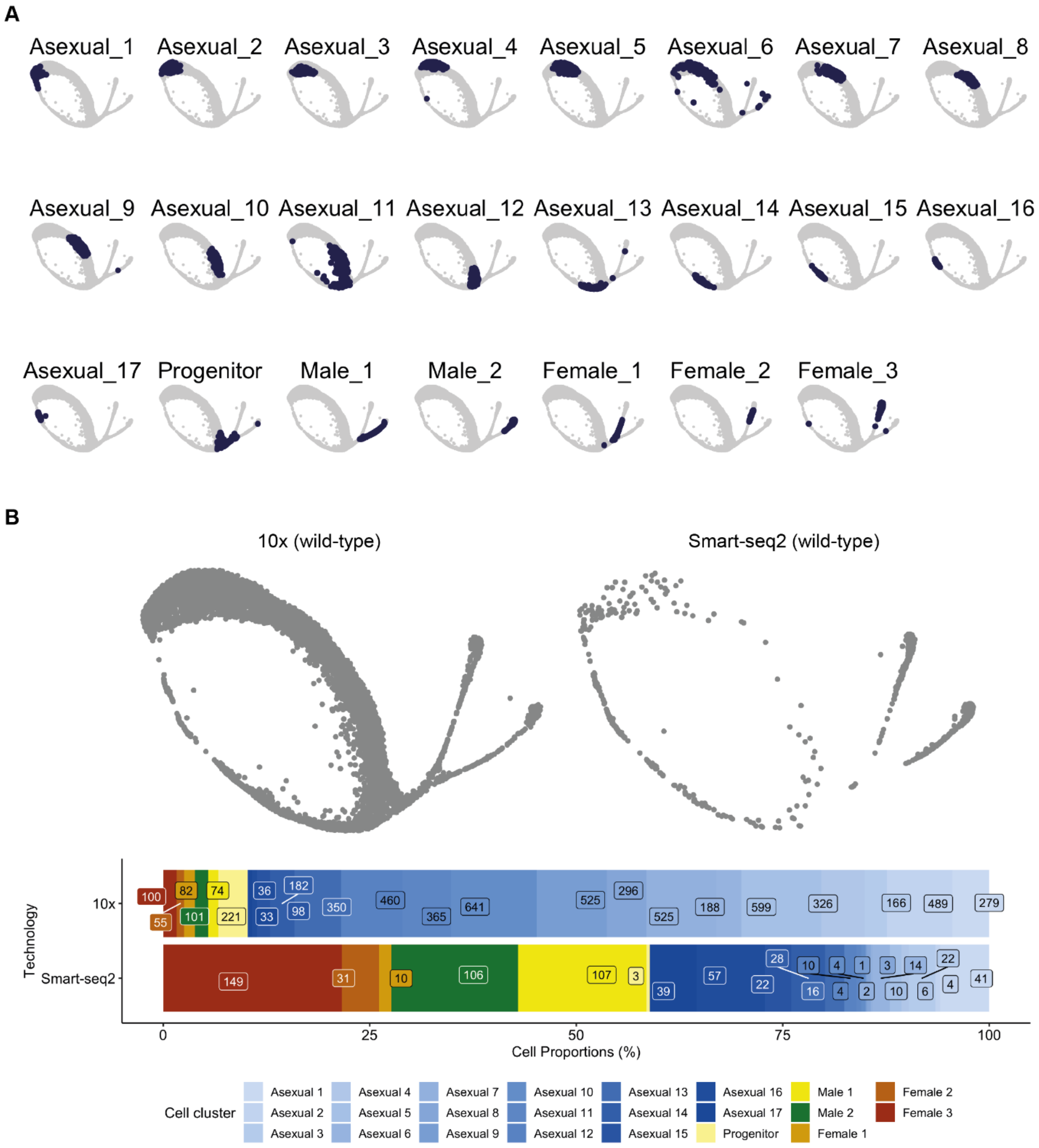


**Fig. S4. Wild-type scRNA-seq Data. (A)** The position of cells on the UMAP for each cluster is shown. **(B)** (top) Deconstructed UMAPs showing contribution of 10x and Smart-seq2 data to the combined analysis. (bottom) bar plot showing the proportion of each cell cluster captured using each technology. Together, these show how Smart-seq2 data captures predominantly later stage parasites, whereas 10x captures all stages.

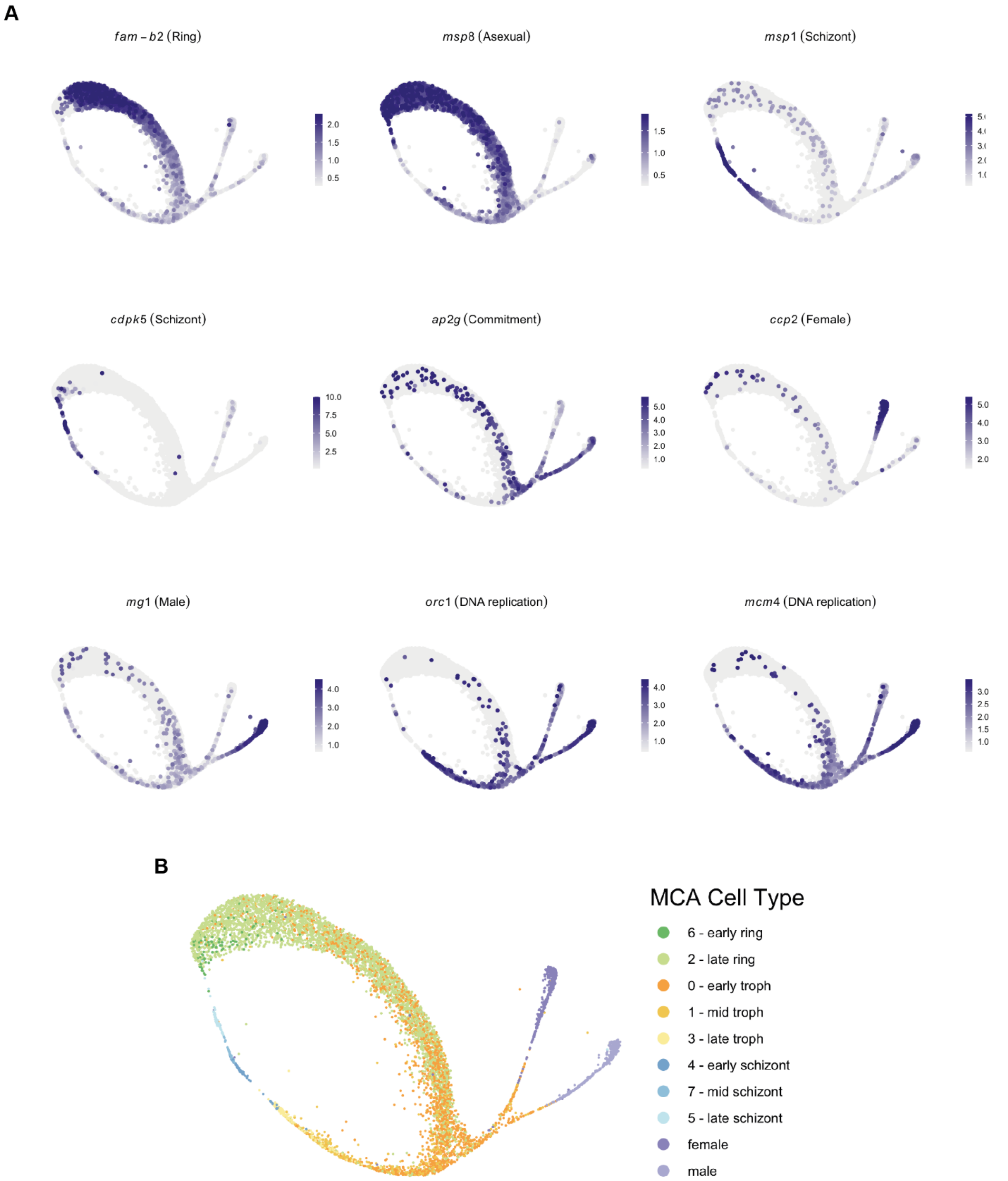


**Fig. S5. Wild-type cell annotation. (A)** The expression of known marker genes in wild-type cells is shown with the 5th and 95th quantiles set as the minimum and maximum expression values respectively to eliminate any outliers having a strong influence on visualisation. Examples include: Fam-b2 (*71*), msp8 (*84*, *85*), msp1 (*86*), CDPK5 (*87*), ap2-g (*3*, *4*), ccp2 (*25*), mg1, orc1 (*88*), mcm4 (*88*). **(B)** Cell types defined by the malaria cell atlas (*89*) were used as a reference to map our complete blood-stage atlas. The original Malaria Cell Atlas did not contain developing gametocytes which is why these cells in the dataset associate most closely to trophozoites.

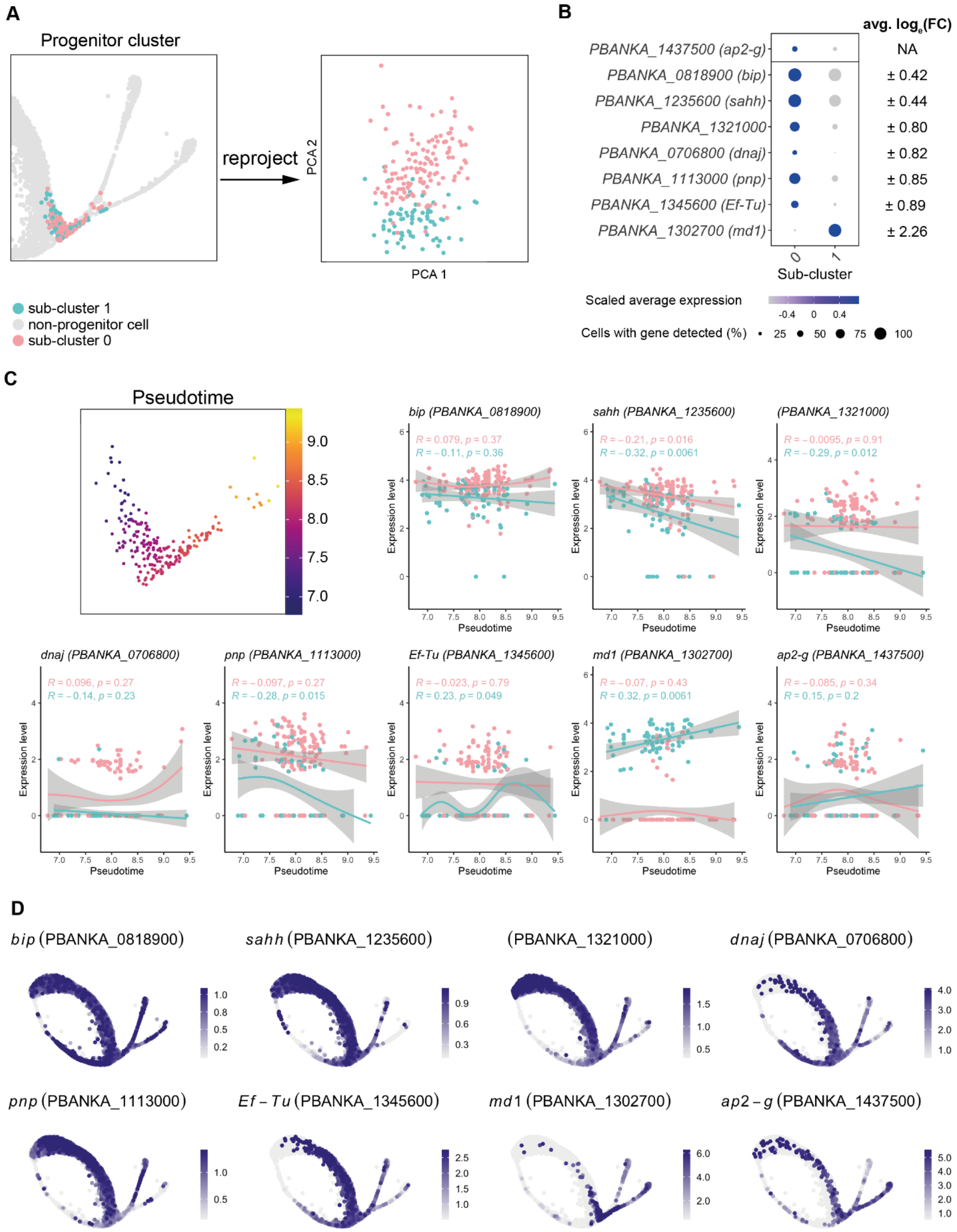


**Fig. S6. Examination of the bipotential branch. (A)** The progenitor cluster was subsetted and re-projected into PCA space. The subset was re-clustered to look for subpopulations (i.e. cells that have committed to one sex over the other) and two subclusters were generated, 0 and 1. **(B)** Marker genes were calculated for each cluster and the expression of statistically-significant (adjusted $p < 0.05$) genes is shown in the dot plot. The only gene with an average $\log_2$(fold-change) > ±1 was *md1*. *ap2-g* is shown for comparative purposes but was not significantly differentially expressed between the two sub-clusters. **(C)** The cells are undergoing a developmental process in the bipotential cluster (left). To ensure that the two sub-clusters did not arise through two phases in progenitor progression over pseudotime, the expression of *md1* and *ap2-g* is plotted against pseudotime (right). Although the expression of *md-1* does increase over pseudotime in sub-cluster 1, the cells of the two subpopulations are dispersed throughout pseudotime. **(D)** To investigate whether the marker genes may have sex-specific roles, their expression is plotted All sub-cluster 0 positive markers show expression in the female branch, with some displaying stark contrast (e.g. *sahh*).

---

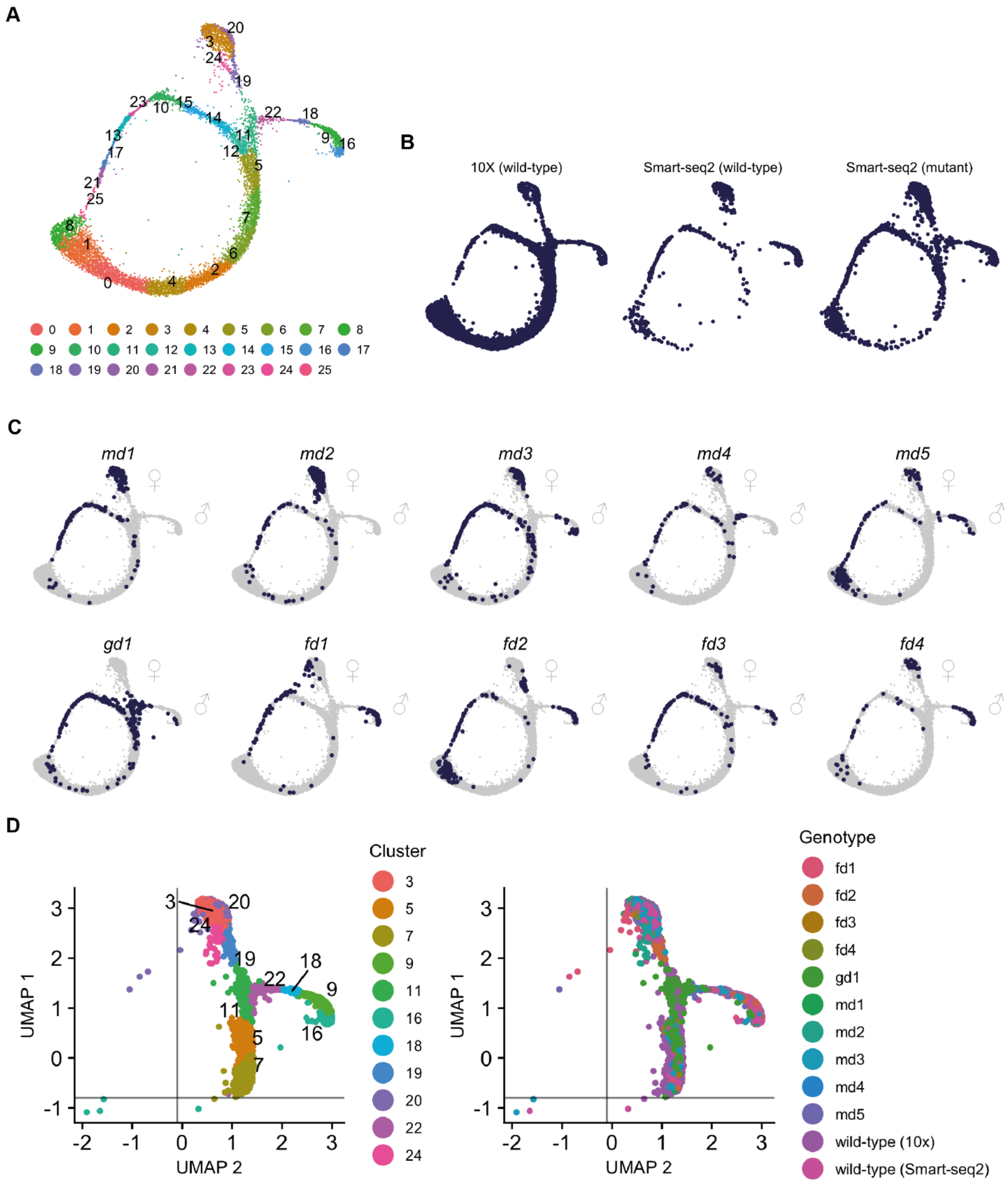

**Fig. S7. Integration of Smart-seq2 and 10x scRNA-seq Data. (A)** UMAP plot of all 8908 cells combined and coloured according to their cluster. **(B)** Deconstructed UMAPs showing contribution of 10x and Smart-seq2 data to the combined analysis. **(C)** UMAP plots highlighting the position of cells belonging to the mutant genotype denoted at the top of the plot. **(D)** UMAP plots showing the cells selected for the sexual branch analysis. Lines show the cutoffs used to remove the 8 outlier cells. Each plot is coloured by cluster membership (left) or genotype (right).

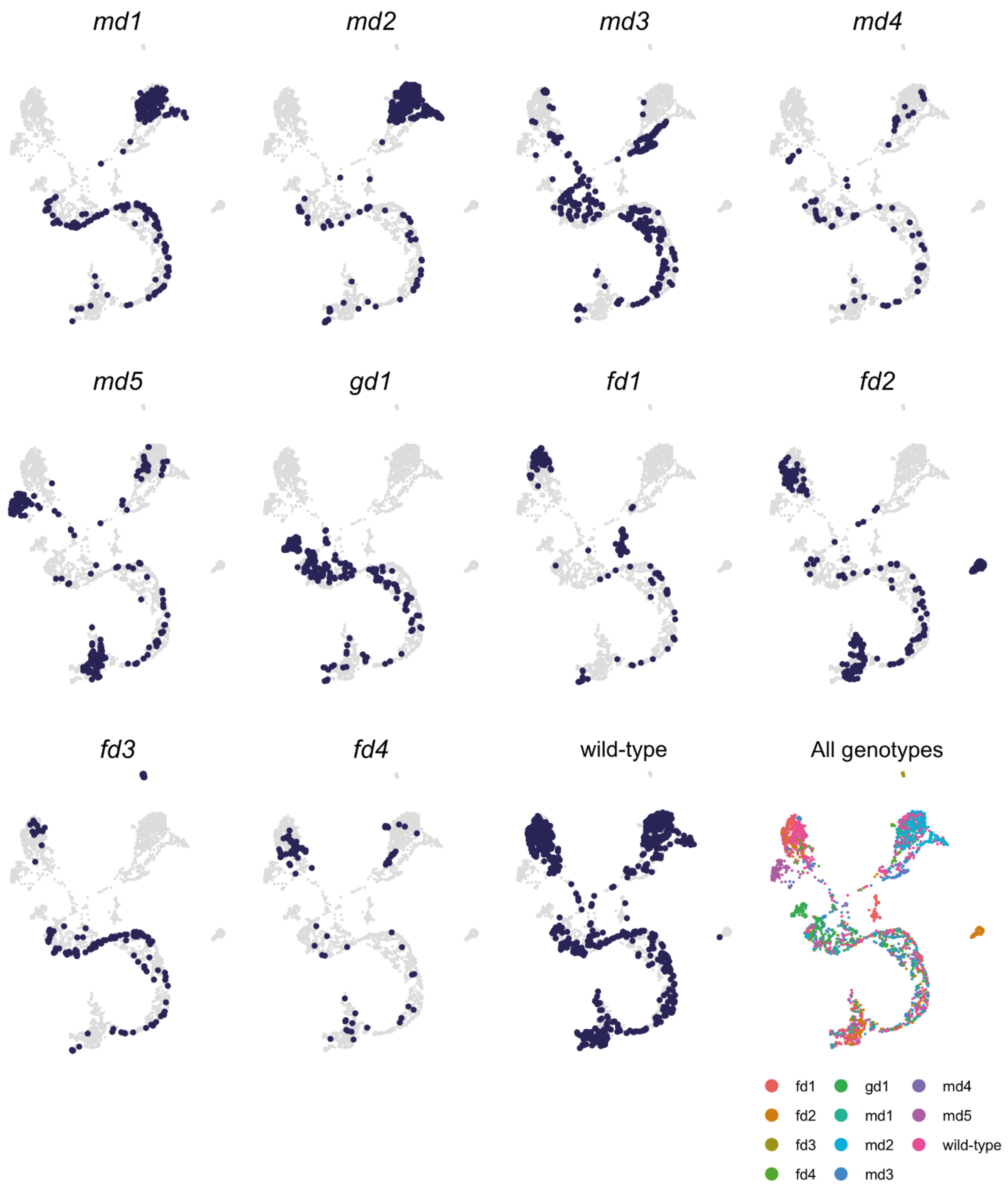


**Fig. S8. Smart-seq2 cells show distinct expression profiles in UMAP plots.** Smart-seq2-only cells were used to generate an alternative UMAP plot. Cells are then coloured according to their genotype as denoted at the top of each plot. This shows that the mutants show distinct transcriptional profiles in the specific sex where they affect development to both each other and mature wild-type male and female gametocytes. This representation can be viewed interactively at: http://obilab.molbiol.umu.se/gcsko/

A

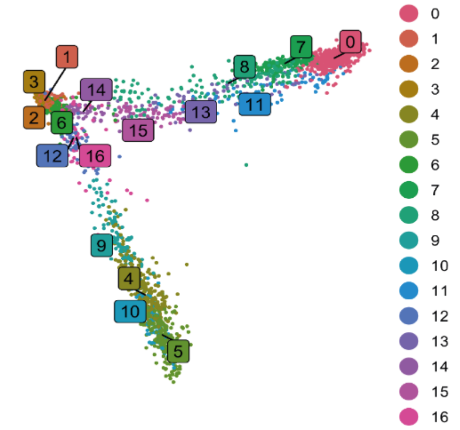


B

Used for DE analysis: mutant (red), wild-type (blue)

| Genotype | 0 | 1 | 2 | 3 | 4 | 5 | 6 | 7 | 8 | 9 | 10 | 11 | 12 | 13 | 14 | 15 | 16 |
|---|---|---|---|---|---|---|---|---|---|---|---|---|---|---|---|---|---|
| fd1 | 0 | 0 | 1 | 1 | 28 | 31 | 0 | 0 | 22 | 2 | 2 | 0 | 0 | 1 | 0 | 0 | 3 |
| fd2 | 0 | 0 | 0 | 1 | 15 | 30 | 2 | 0 | 2 | 3 | 6 | 0 | 0 | 0 | 3 | 57 | 0 |
| fd3 | 9 | 5 | 1 | 2 | 7 | 5 | 0 | 5 | 0 | 2 | 0 | 5 | 0 | 0 | 0 | 0 | 0 |
| fd4 | 11 | 0 | 1 | 0 | 24 | 14 | 0 | 9 | 2 | 4 | 3 | 1 | 0 | 0 | 0 | 0 | 0 |
| gd1 | 0 | 17 | 4 | 8 | 0 | 0 | 14 | 0 | 0 | 0 | 0 | 0 | 25 | 0 | 1 | 0 | 29 |
| md1 | 66 | 4 | 2 | 3 | 0 | 0 | 0 | 42 | 4 | 0 | 0 | 4 | 0 | 11 | 0 | 0 | 1 |
| md2 | 102 | 3 | 0 | 1 | 0 | 0 | 0 | 55 | 1 | 0 | 0 | 4 | 0 | 42 | 0 | 1 | 0 |
| md3 | 19 | 10 | 5 | 6 | 28 | 2 | 2 | 6 | 50 | 2 | 2 | 11 | 1 | 1 | 0 | 0 | 9 |
| md4 | 7 | 1 | 2 | 1 | 0 | 0 | 1 | 7 | 0 | 0 | 0 | 1 | 16 | 0 | 0 | 0 | 0 |
| md5 | 15 | 0 | 0 | 0 | 1 | 0 | 2 | 1 | 4 | 58 | 45 | 4 | 2 | 0 | 0 | 0 | 4 |
| wild-type (10x) | 80 | 325 | 341 | 266 | 42 | 42 | 169 | 6 | 31 | 31 | 20 | 36 | 34 | 1 | 65 | 3 | 2 |
| wild-type (Smart-seq2) | 84 | 4 | 2 | 6 | 118 | 73 | 4 | 54 | 24 | 9 | 10 | 15 | 1 | 18 | 0 | 1 | 5 |

Cluster

C

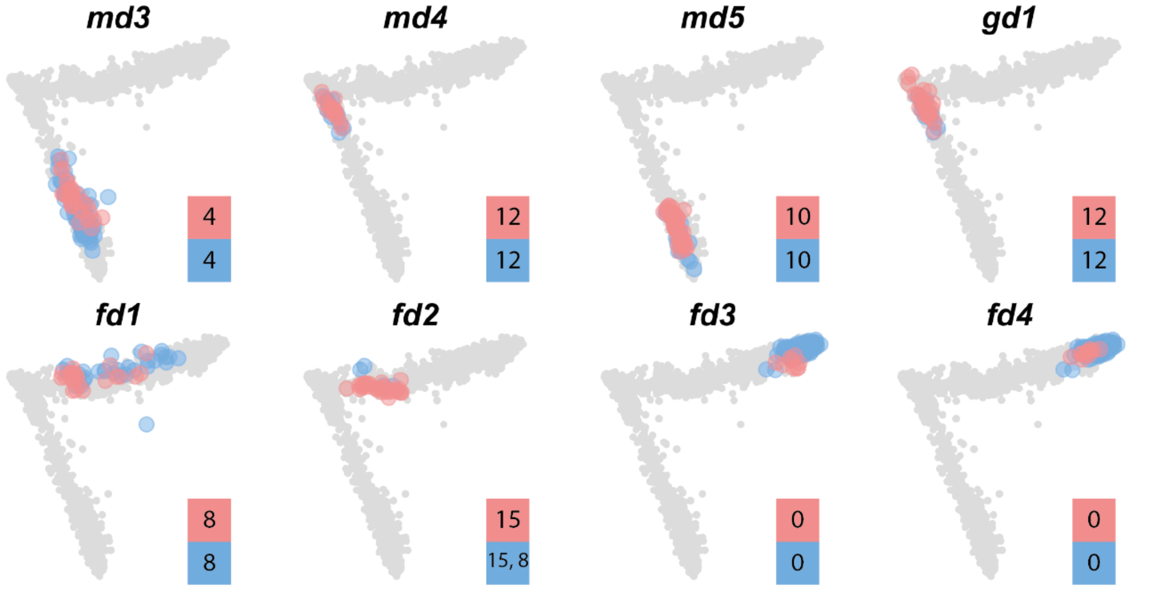


**Fig. S9. Differential expression testing. (A)** PCA plot showing the clusters that were generated after subsetting the branch cells in the wild type and mutant combined dataset. **(B)** Table showing the clusters that were used for differential expression analysis for each mutant (red) and the corresponding wild-type cells that were tested against (blue). The number of cells in each class is shown in the box. **(C)** PCA plots showing where the clusters that were used for differential expression analysis are located. The cluster number used is denoted in the boxes, top = mutant cell cluster, bottom = wild-type cell cluster. Using clusters that are similarly located on the PCA reduces the possibility that DE genes identified are due to differences in progression along the branch caused by the mutation rather than the mutation itself.

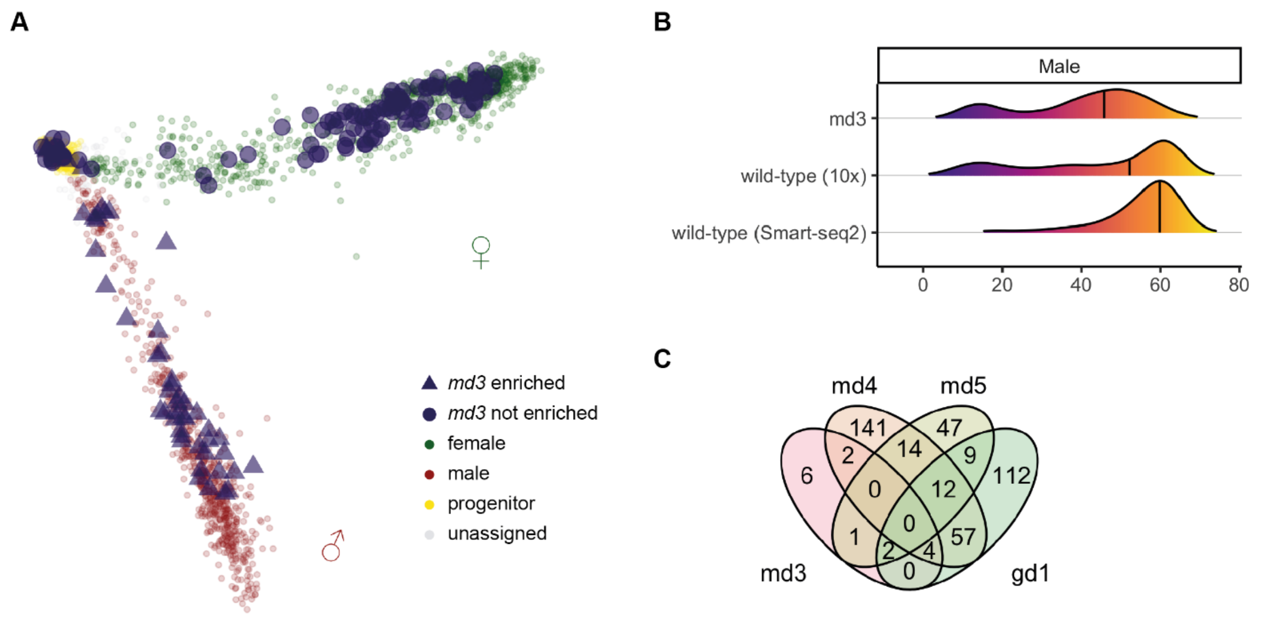


**Fig. S10. Targeted analysis of *md3* male gametocytes.** Disruption of *md3* was associated with a loss of male marker expression in the screen, and all 87 gametocytes subjected to Smart-seq2 analysis were females, indicative of a shift in sex ratio. However, in contrast to *md1* and *md2* mutants, male fertility was not completely lost in a genetic cross with a *hap2* mutant (Fig. 2F), and an *md3* KO clone was able to transmit the disrupted *md3* locus to mosquitoes and back to mice (Fig. 2E, see Fig. S8 for genotype confirmation of the transmitted parasites), suggesting there was a strongly reduced number of males in this mutant, which remained fertile. We therefore enriched rare cells expressing the male reporter gene by flow cytometry and subjected these to Smart-seq2 analysis. (**A**) Smart-seq2 transcriptomes of enriched *md3* male gametocytes mapped onto the PCA plot from Fig. 4A. (**B**) Frequency distribution of transcriptomes over pseudotime for male md3 gametocytes enriched by flow sorting and wild type for each technology. (**C**) Numbers of genes with differences in transcript abundance in male gametocytes, compared to wild type as in Fig. 4C, but including the combined transcriptomes of male *md3* gametocytes enriched by flow sorting.

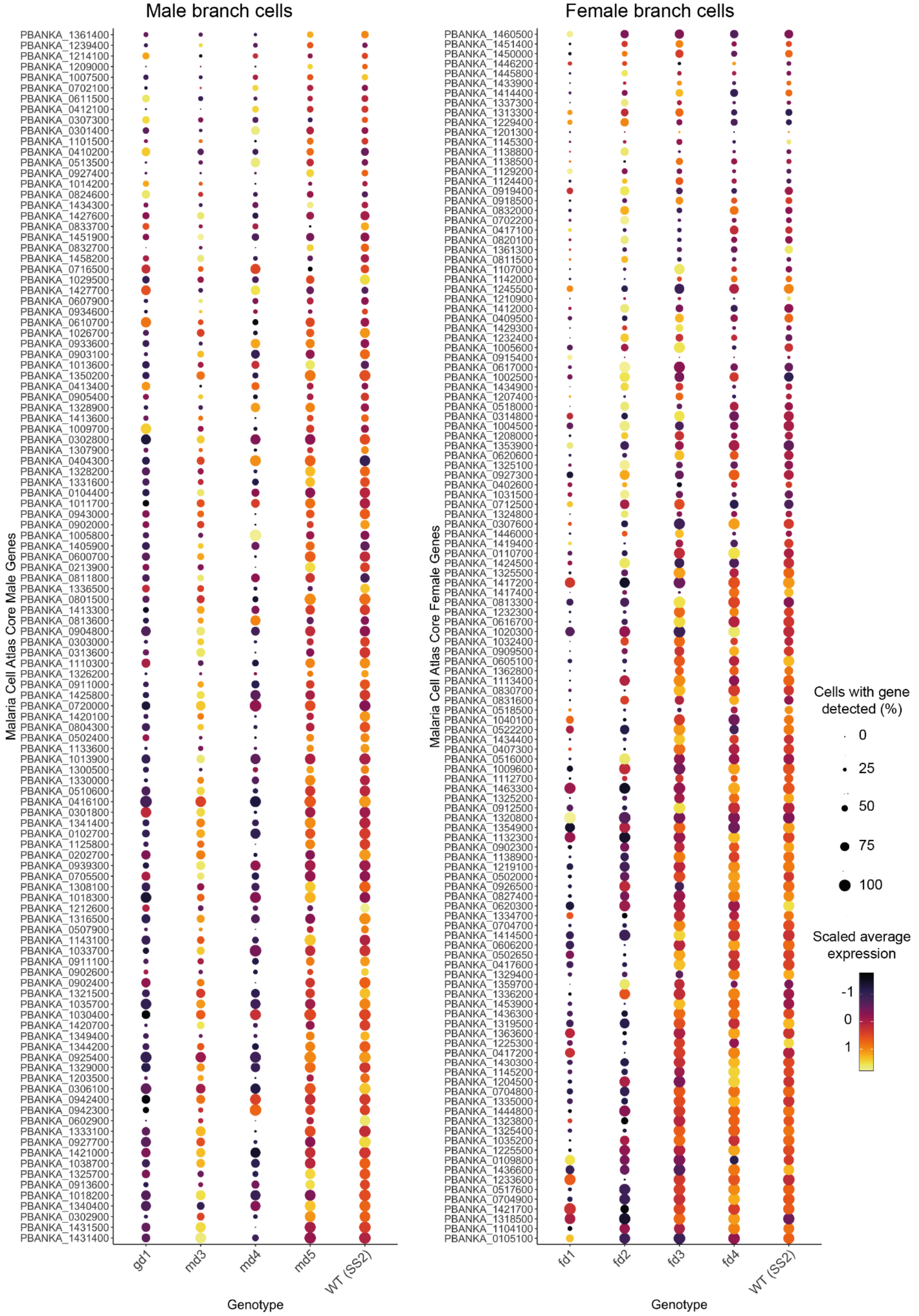


**Fig. S11. Core Sexual Genes Analysis.** The expression of core sexual genes defined in (*89*) were plotted in each genotype within each sexual branch. WT (SS2) = wild-type (Smart-seq2).

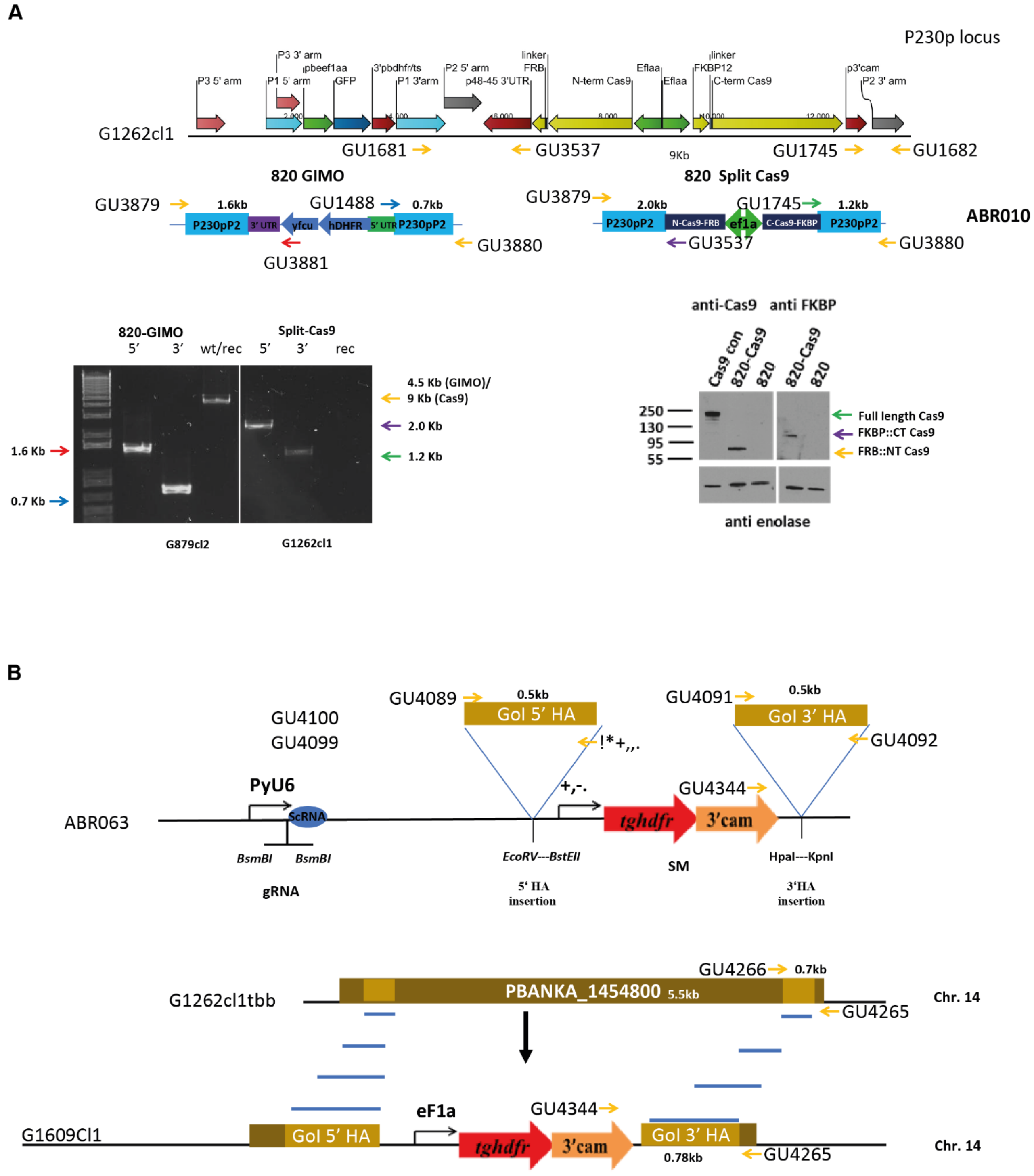


**Fig. S12. Strategy for the inducible targeted disruption of *fd1* using a split Cas9 recombinase. (A)** Top line: Modified p230p locus after insertion of an expression cassette for an carboxy-terminal Cas9 fragment fused to FKBP and an amino-terminal fragment fused to FRB, rendering Cas9 inducible by addition of rapamycin analogues. The two-step approach involved the initial insertion of a postive-negative selection cassette (bottom line, left) under selection for a human dhfr gene next to an expression cassette for GFP, followed by the replacement of the selection cassette with the slip Cas9 cassette under negative selection against the yfcu gene. **(B)** PCR products of the expected size confirm the successful insertion of split Cas9 into the redundant *p230p* locus. **(C)** Western blot analysis showing expression of N-terminal Cas9 using an antibody directed against Cas9. Expression of the C-terminal Cas9 fragment is demonstrated using an antibody against the fusion partner FKBP. **(D)** Schematic illustrating the CRISPR-Cas9-mediated disruption of *fd1* using targeting plasmid ABR063 to replace most of the *fd1* open reading frame.

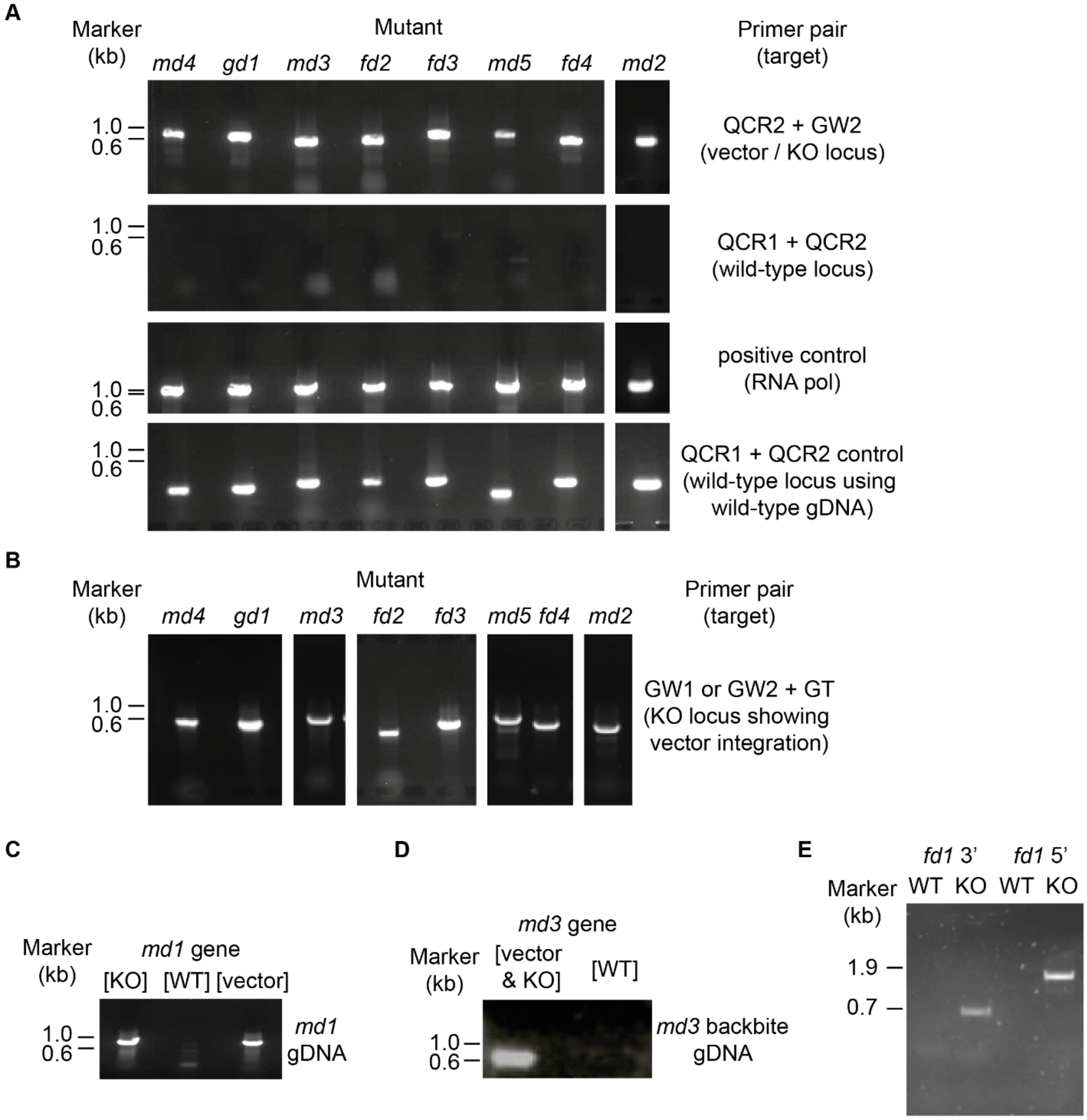


**Fig. S13. PCR products confirm knockout genotypes of cloned mutants. (A)** PCR products showing presence of disrupted and absence of wild type loci on mutant DNA in mutants made with *Plasmo*GEM vectors (lower panel control gDNA from wild type). **(B)** Additional, longer PCR products showing genomic integration of the targeting vectors. **(C)** Genotype evidence for the *md1* mutant. **(D)** PCR evidence showing *md3* mutant parasites which have passed through mosquitoes retained the disrupted allele. **(E)** PCR verification of 3' and 5' CRISPR-induced integration of a deletion vector for *fd1*.

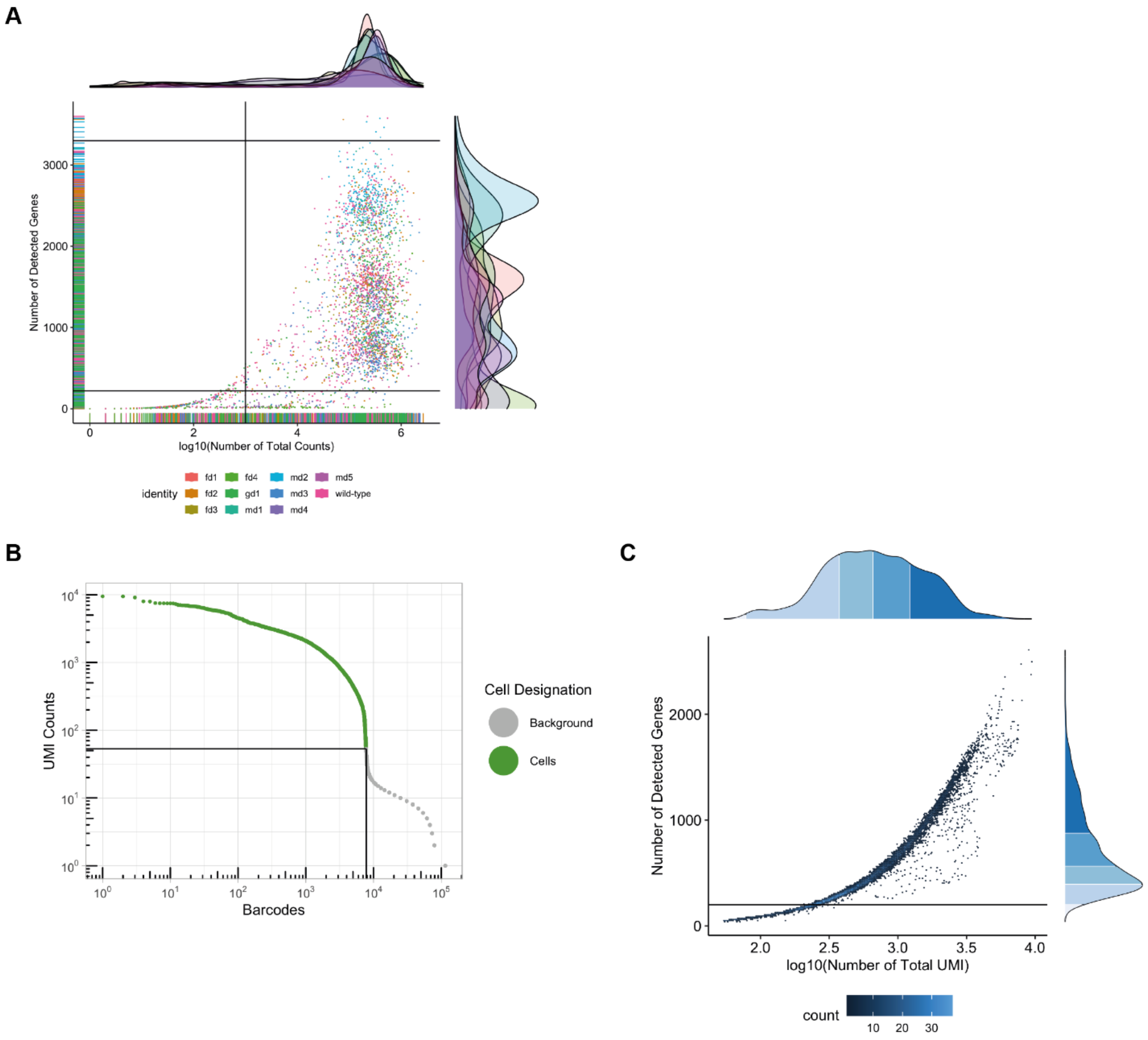


**Fig. S14. Quality control of single-cell RNA-seq data. (A)** A plot showing all Smart-seq2 cells before quality control and the thresholds used to eliminate low-quality cells. Cells are coloured according to their genotype. **(B)** A plot of the 10x data showing the number of UMI counts detected per cell and the cumulative number of barcodes. This distribution was used to define high-quality cells vs. background. **(C)** A plot of the 10x data after definition of the cells, showing the number of detected genes per cell threshold used in quality control.
